# CellO: Comprehensive and hierarchical cell type classification of human cells with the Cell Ontology

**DOI:** 10.1101/634097

**Authors:** Matthew N. Bernstein, Zhongjie Ma, Michael Gleicher, Colin N. Dewey

## Abstract

Cell type annotation is a fundamental task in the analysis of single-cell RNA-sequencing data. In this work, we present CellO, a machine learning-based tool for annotating human RNA-seq data with the Cell Ontology. CellO enables accurate and standardized cell type classification by considering the rich hierarchical structure of known cell types, a source of prior knowledge that is not utilized by existing methods. Furthemore, CellO comes pre-trained on a novel, comprehensive dataset of human, healthy, untreated primary samples in the Sequence Read Archive, which to the best of our knowledge, is the most diverse curated collection of primary cell data to date. CellO’s comprehensive training set enables it to run out-of-the-box on diverse cell types and achieves superior or competitive performance when compared to existing state-of-the-art methods. Lastly, CellO’s linear models are easily interpreted, thereby enabling exploration of cell type-specific expression signatures across the ontology. To this end, we also present the CellO Viewer: a web application for exploring CellO’s models across the ontology.

**Highlight:**

- We present CellO, a tool for hierarchically classifying cell type from single-cell RNA-seq data against the graph-structured Cell Ontology
- CellO is pre-trained on a comprehensive dataset comprising nearly all bulk RNA-seq primary cell samples in the Sequence Read Archive
- CellO achieves superior or comparable performance with existing methods while featuring a more comprehensive pre-packaged training set
- CellO is built with easily interpretable models which we expose through a novel web application, the CellO Viewer, for exploring cell type-specific signatures across the Cell Ontology

**Graphical Abstract:** 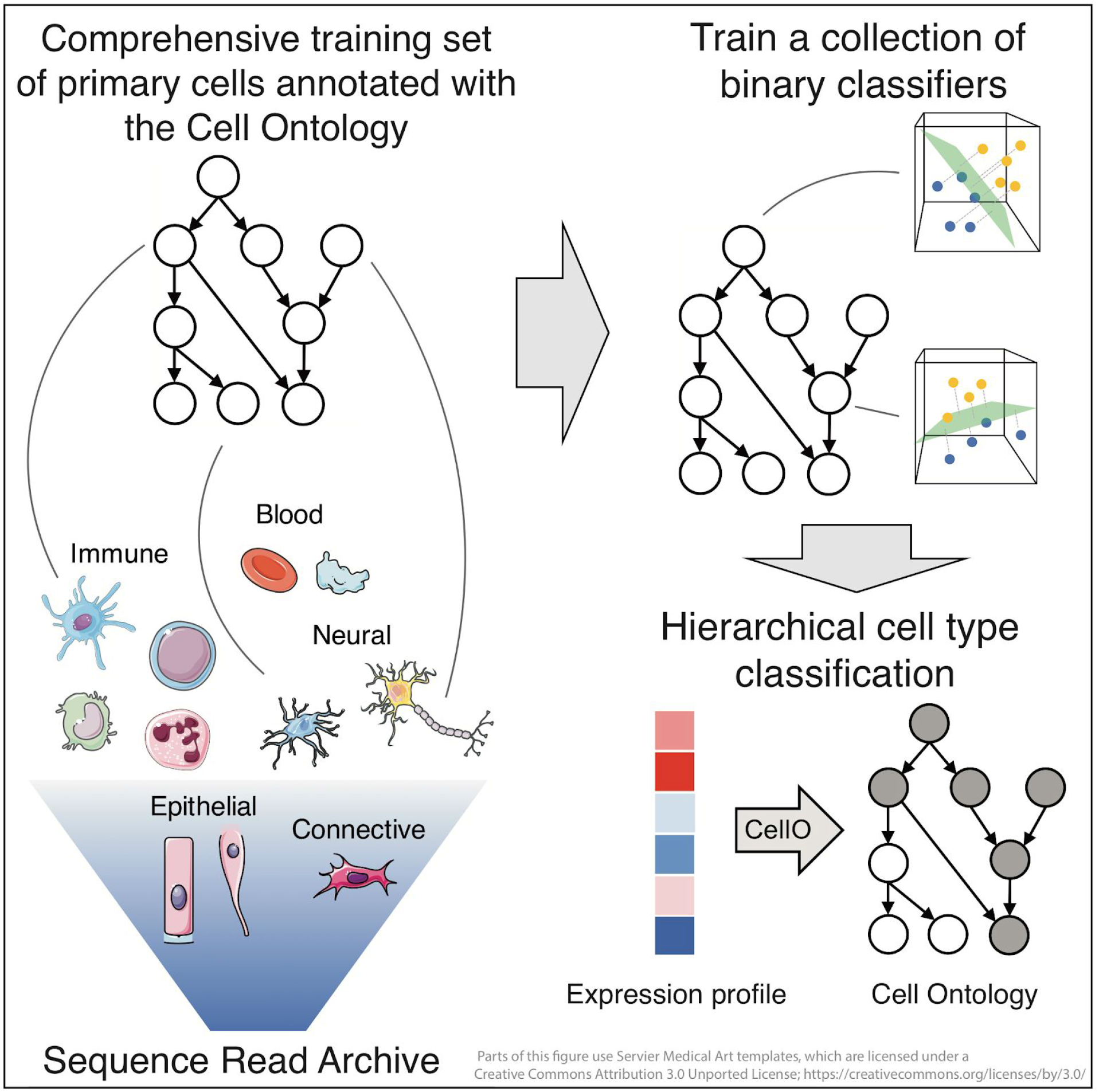

## Introduction

Cell type annotation is a fundamental task in the analysis of single-cell RNA-sequencing (scRNA-seq) data. Recently, a number of computational tools have been developed for automating the cell type annotation task. Nonetheless, many of these tools suffer from certain disadvantages that inhibit their use. First, many existing methods require the user to provide either a set of marker genes associated with each cell type (Pliner et al., 2019; Zhang et al., 2019a) or a suitable training dataset with cells already annotated with cell type labels (Alquicira-Hernandez et al., 2019; Ma and Pellegrini, 2020; Tan and Cahan, 2019). Marker gene-based approaches are challenged by the fact that there is not a canonical set of marker genes for most cell types (Zhang et al., 2019b). Furthermore, finding an appropriate and labelled training set that contains all of the cell types in the target dataset can be challenging, especially considering that existing approaches are sensitive to the chosen training set (Abdelaal et al., 2019).

Second, many existing methods use flat-classification. Flat classification suffers from the possibility that predictions are logically inconsistent with the hierarchy of cell types. Specifically, for a given query, a flat classifier may output a probability for a cell type that is larger than the classifier’s output for its parent cell type in the hierarchy (Obozinski et al., 2008). Such incoherent outputs reduce the scientific usefulness of the classifier. We assert that framing the cell type classification task as that of hierarchical classification against the Cell Ontology (Bard et al., 2005) poses a number of advantages over flat-classification. The Cell Ontology provides a comprehensive hierarchy of animal cell types encoded as a directed acyclic graph (DAG). This DAG provides a rich source of prior knowledge to the cell type classification task that remains un-utilized in flat classification. In addition, if the algorithm is uncertain about which specific cell type the cell may be, the use of a hierarchy allows the algorithm to place a cell internally within the graph rather than at a leaf node.

Finally, those methods that do perform hierarchical classification do not make use of the rich hierarchical relationships between known cell types encoded by the Cell Ontology. For example, CHETAH (de Kanter et al., 2019) classifies cells against a hierarchy; however, CHETAH infers this hierarchy from the data rather than utilizing the existing hierarchy encoded by the Cell Ontology. Garnett (Pliner et al., 2019) utilizes a hierarchy of cell types; however, these hierarchies must be pre-specified by the user. Furthermore, Garnett requires that each cell within the hierarchy be associated with a set of marker genes. To the best of our knowledge, the only method that utilizes the graph-structure of an ontology for the task at hand is URSA (Lee et al., 2013), which classifies gene expression profiles against the BRENDA Tissue Ontology (Gremse et al., 2011).

In this work, we present Cell Ontology-based Classification (CellO) a tool for annotating cells against the graph-structured Cell Ontology (**Figure 1A**). CellO is a discriminative, supervised machine learning approach that comes pre-trained on a novel comprehensive dataset comprising nearly all human primary samples in the Sequence Read Archive (SRA; Leinonen et al., 2011) and therefore arrives ready-to-run on diverse scRNA-seq datasets. Lastly, CellO makes extensive use of linear models, which are particularly amenable to interpretation. To enable their interpretation, we present a web-based tool, the CellO Viewer, for exploring the cell type expression signals uncovered by the model (https://uwgraphics.github.io/CellOViewer/). We benchmarked CellO on a collection of diverse single-cell datasets and found CellO capable of accurately annotating datasets that existing state-of-the-art and ready-to-run (i.e. come pre-trained) annotation methods were unable to accurately annotate, thus highlighting CellO’s ability to annotate diverse datasets out-of-the-box. Through its use of the Cell Ontology, its comprehensive training set, and the interpretability of its models, CellO addresses the aforementioned limitations of existing tools, thus providing a practical tool for scRNA-seq cell type annotation. (**Figure 1B**).

**Figure 1.**
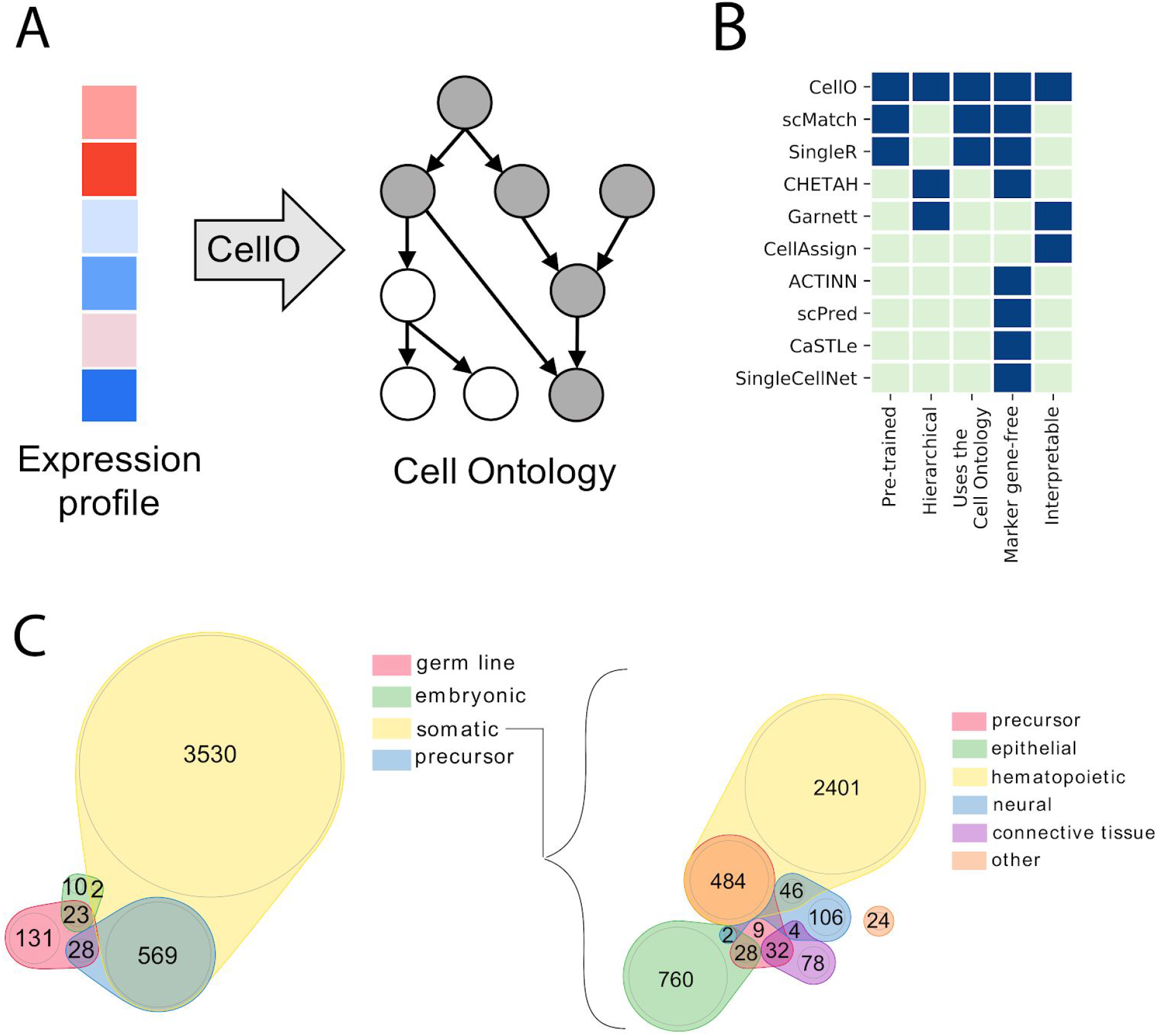
Overview of CellO. (**a**) A schematic overview of CellO’s hierarchical classification approach. CellO performs hierarchical classification with the Cell Ontology. Given a gene expression profile, CellO annotates the cell with a set of cell types (grey nodes) that are consistent with the hierarchical structure of the Cell Ontology. (**b**) We compare CellO to eight recent cell type annotation methods regarding the criteria we surmise are desirable in a cell type classification approach: whether the method (1) arrives pre-trained and can run out-of-the-box, (2) incorporates a hierarchy of cell types, (3) specifically uses the Cell Ontology as its hierarchy, (4) requires cell type-specific marker genes, and (5) uses models can be interrogated to better understand how they arrived at their decision. We compare CellO to scMatch (Hou et al., 2019), SingleR (Aran et al., 2019), CHETAH (de Kanter et al., 2019), Garnett (Pliner et al., 2019), CellAssign (Zhang et al., 2019a), ACTINN (Ma and Pellegrini, 2020), scPred (Alquicira-Hernandez et al., 2019), CaSTLe (Lieberman et al., 2018), and SingleCellNet (Tan and Cahan, 2019). CellO meets all desirable criteria while existing approaches fail to meet at least one. (**c**) Euler diagrams of the cell types within the bulk RNA-seq expression profiles used to train CellO. This training set comprises most primary cell bulk RNA-seq samples within the SRA and consists of diverse cell types spanning various tissues, developmental stages, and stages of differentiation. These diagrams were created with nVenn (Pérez-Silva et al., 2018).

## Results

### A novel curated RNA-seq dataset of human primary cells

In order to capture robust cell type signals, we sought a dataset of bulk RNA-seq samples comprising only healthy primary cells that originate from cells that have been isolated based on phenotypic characteristics downstream of gene expression itself (such as cell surface proteins). We thus avoid the circularity in using ground truth cell type labels determined by gene expression (via the expression of cell type-specific marker genes) as are often provided in scRNA-seq datasets. We did not wish to include cells that underwent multiple passages, were diseased, or underwent other treatments, such as in vitro differentiation, because these conditions alter gene expression. We therefore curated a novel dataset from the SRA consisting of healthy, untreated, primary cells. To do so efficiently, we leveraged the annotations provided by the MetaSRA project (Bernstein et al., 2017) which includes sample-specific information including cell type, disease-state, and sample type. We then manually curated the samples selected via the MetaSRA by both annotating technical variables and refining cell type annotations (Methods).

This curation effort resulted in a dataset comprising 4,293 bulk RNA-seq samples from 264 studies. These samples were labeled with 310 cell type terms, of which 113 were the most-specific cell types in our dataset (i.e., no sample in our data was labelled with a descendent cell type term). These cell types were diverse, spanning multiple stages of development and differentiation (**Figure 1c**). We uniformly quantified and normalized (via log transcripts per million) gene expression from the raw RNA-seq data for these samples (Methods). To the best of our knowledge, this dataset is the largest and most diverse set of bulk RNA-seq samples derived from only primary cells. Prior to this work, the most comprehensive bulk primary cell transcriptomic dataset was compiled by (Aran et al., 2017), which contains data for 64 cell types from 6 studies. Whereas our dataset consists of only RNA-seq data, this prior dataset included samples assayed with several other technologies, such as microarrays. Another comprehensive set of primary cell expression data was collected by (Mabbott et al., 2013), which contains primary cell data from 745 samples from 105 studies; however, these data are exclusively from microarrays.

### Novel applications of hierarchical classification methods

We frame the cell type classification task as hierarchical classification against the Cell Ontology’s DAG. The hierarchical classification task is inherently a multi-label classification task where each input sample (i.e. cell) is mapped to a *set* of output labels (i.e. cell types). Hierarchical classification extends multi-label classification by further requiring that the output labels are *consistent* with the labels’ DAG. That is, for each label in a given output set of labels, the label’s parent labels are also in the output set (**Figure 1A**).

We implemented two strategies for performing hierarchical classification against the Cell Ontology’s DAG that both come packaged with CellO. First, we implemented cascaded logistic regression (CLR; Obozinski et al., 2008), which entails classifying a sample in a top-down fashion from the root of the ontology downward via a collection of binary classifiers. Specifically, each binary classifier is associated with a cell type and is trained to classify a sample conditioned on the sample belonging to all of the cell type’s parents in the ontology.

Next, we implemented a collection of one-vs.-rest binary classifiers for each cell type in the DAG. We will refer to this as the “independent classifiers” approach. This approach suffers from the possibility that the classifiers’ outputs will be inconsistent with the hierarchical structure of the ontology. An inconsistency occurs when the output probability for a given cell type exceeds that of one of its parent cell types in the ontology. We tested the use of independent logistic regression classifiers and found inconsistencies to be a frequent source of errors. Specifically, we performed leave-study-out cross-validation on the full set of bulk RNA-seq data and examined the consistency of all edges that were adjacent to at least one cell type whose classifier produced a non-negligible probability (> 0.01) of the sample originating from that cell type. Of these edges, 12.1% were inconsistent (**Figure S1**). Nearly all samples (>99%) contained at least one inconsistent edge and 34% of samples contained at least one severely inconsistent edge in which the child classifier’s probability exceeded the parent classifier’s probability by at least 0.25. We will use the term “correction” to refer to the task of reconciling the outputs of independent classifiers with a hierarchy (**Figure 2A**).

**Figure 2.**
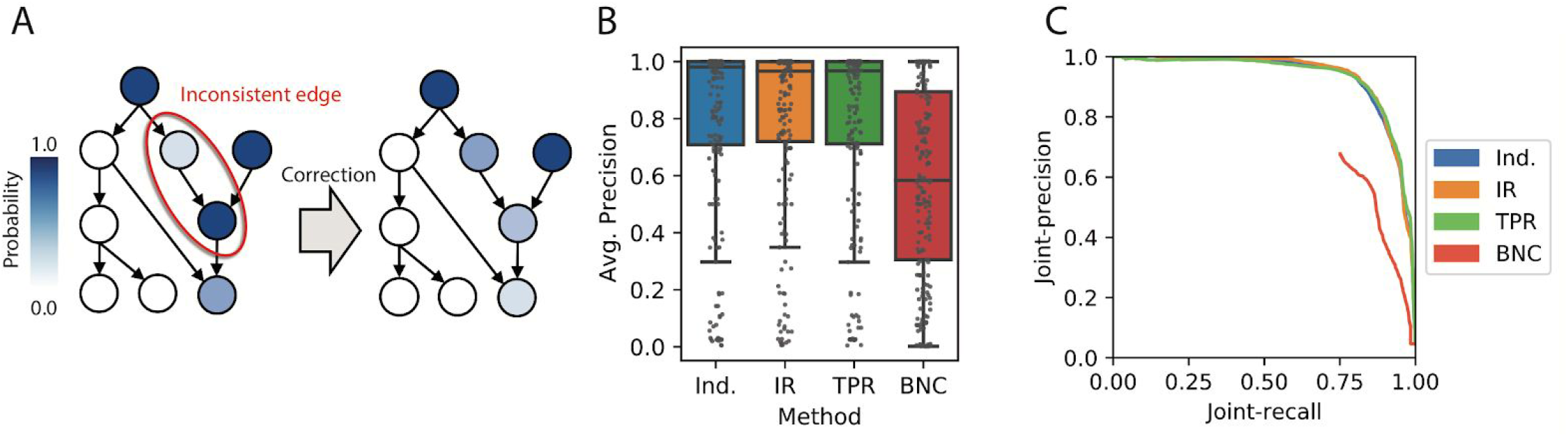
Reconciling the outputs of independent classifiers with a hierarchy. (**A**) A schematic illustration of the reconciliation task for correcting the outputs of independent classifiers according to a hierarchy. The left DAG’s nodes are colored according to the raw classifier probabilities for each node. Highlighted is one edge of the graph whose incident nodes have probabilities that are logically inconsistent with the hierarchy because the child node has a higher probability than the parent. The correction task involves modifying these probabilities such that all probabilities are consistent with the hierarchy (right). (**B**) F1-scores across all cell types for the independent classifiers (Ind.), as well as for IR, TPR, and BNC on the validation set. (**C**) Each paired sample and cell type prediction was considered independently. The set of all such predictions was ordered according to their prediction probability and the corresponding precision-recall curve was constructed for the independent classifiers, IR, TPR, and BNC.

To date, the one correction method that has been applied to the task at hand is Bayesian network correction (BNC), which is implemented in the URSA tool (Lee et al., 2013). Therefore, as a baseline, we implemented a BNC algorithm following the description in Lee *et al.* (2013) (Methods). We also tested two correction methods that have yet to be applied to the cell type classification task: isotonic regression correction (IR; Obozinski et al., 2008) and a heuristic procedure called the True Path Rule (TPR; Notaro et al., 2017). IR uses a projection-based approach for correction that entails finding a set of consistent output cell type probabilities that minimize the sum of squared differences to the raw, and possibly inconsistent, classifier output probabilities. In contrast, TPR uses a heuristic procedure that involves a bottom-up pass through the ontology such that the output of children classifiers are averaged with the output of the parent classifier to allow information flow across the ontology graph.

To test these correction methods, we first partitioned the bulk RNA-seq dataset into a pre-training and validation set (**Figure S4**; Methods). Using this validation set, we performed a grid search to find the optimal parameters for training each binary logistic regression classifier, and given the optimal set of parameters, compared how well the aforementioned correction methods either enhanced or degraded accuracy over the samples in the validation set. Overall, we find that IR and TPR output probabilities similar to those output by the independent classifiers in regards to both average precision scores across the cell types in the validation set (**Figure 2B**) and precision-recall curves when considering each sample-cell type pair as an independent prediction (**Figure 2C**). This indicates that IR and TPR do not degrade performance in comparison to independent classifiers. In contrast, we found that the BNC approach significantly degraded performance (**Figure 2B, C**). We note that these results are in line with work by Obozinski *et al* (2008), which demonstrates that IR outperforms BNC on the hierarchical protein function prediction task. Although both IR and TPR yielded similar results, we use IR as our correction method of choice due to its simplicity.

We also used this partition of the bulk RNA-seq samples to tune the parameters of the CLR algorithm (Methods). We found that after tuning, both IR and CLR achieved similar maximum median F1-scores and median average-precisions across cell types on the validation set (Figure S5) and therefore both are included in the CellO software and evaluated throughout the remainder of this study

### Evaluation on scRNA-seq data

We trained both CLR and IR on the full set of bulk RNA-seq samples in order to test their performance on single-cell data. We note that training a single-cell classifier with bulk RNA-seq data may lead to models being poorly calibrated to the sparse single-cell expression profiles. To address this challenge, we first cluster single-cell data using the Leiden community-detection algorithm (Traag et al., 2019) using the default resolution parameter of 1.0, as implemented in the Scanpy Python package (Wolf et al., 2018), and then compute each cluster’s mean expression profile. The mean expression profiles are less sparse than those of the individual cells and thus better resemble the bulk RNA-seq data on which the algorithms were trained. CellO first classifies each cluster based on its mean expression profile and then assigns each cell to its cluster’s assigned cell types.

We compiled a dataset consisting of 7,366 healthy primary cells originating from non-droplet-based RNA-seq assays, such as SMART-Seq2 (Picelli et al., 2013) and MARS-seq (Jaitin et al., 2014), from the SRA in a manner similar to that used for compiling the bulk RNA-seq training data. This dataset originated from 14 studies and were labeled with 125 cell type terms, of which 32 were most specific to the data. Of these cells, 4,936 were of cell types that were included in the bulk RNA-seq training set. This subset of cells originate from 12 studies and were labeled with 71 cell type terms of which 16 were most specific to the dataset. We note that for many of the cells used in this analysis, the ground-truth cell types provided by the authors of the data were determined via *in silico* and/or manual approaches (e.g. via heuristic marker-gene based approaches) and thus, this analysis can be understood as an analysis of the *consistency* between the cell types as annotated by the authors and those annotated by the automated methods explored in this work.

We use the subset of 4,936 cells to evaluate IR, CLR, as well as a baseline one-nearest-neighbor (1NN) algorithm that simply returns the cell type labels of the most similar sample in the training set to the query expression profile using Pearson correlation as the similarity metric. We evaluated two aspects of these algorithms’ classifications. First, we compute the average-precision (a measure of area under the precision-recall curve) on each cell type’s output probabilities. Second, for each cell, we evaluate a set of binary yes-no decisions for each cell type that result from thresholding the raw output probabilities and enforce each cell to be annotated with only one most-specific cell type (Methods; **Figure S8**). We evaluate these binary decisions using precision, recall, and F1-score (harmonic mean of precision and recall). We modified the evaluation metrics to take into account samples that were labelled with a general cell type, but not a specific cell type (e.g. T cell versus CD8+ T cell; Methods).

We found that IR and CLR out-performed 1NN according to F1-score, precision, and average-precision (**Figure 3A, B, C**). Specifically, IR, CLR, and 1NN produced median F1-scores of 0.81, 0.85, and 0.63 respectively across all cell types. The cell types on which classification performance was poor were generally more-specific and clustered within the hierarchy (**Figure 3D**). We note that CellO’s average precision scores across cell types tend to be higher than its F1-scores (**Figure 3D**). This discrepancy indicates that CellO is doing well at discriminating among these cell types; however, the decision thresholds used by CellO to output hard-classifications for some cell types may be non-optimal. We hypothesize that this is due to some classifiers being poorly calibrated, especially for cell types lower in the ontology. This may due to there being fewer *studies* in the training set generating these cell types and thus, for a given cell type, CellO’s binary classifiers may be more prone to fitting the batch effects present in these fewer studies (**Figure S9**).

**Figure 3.**
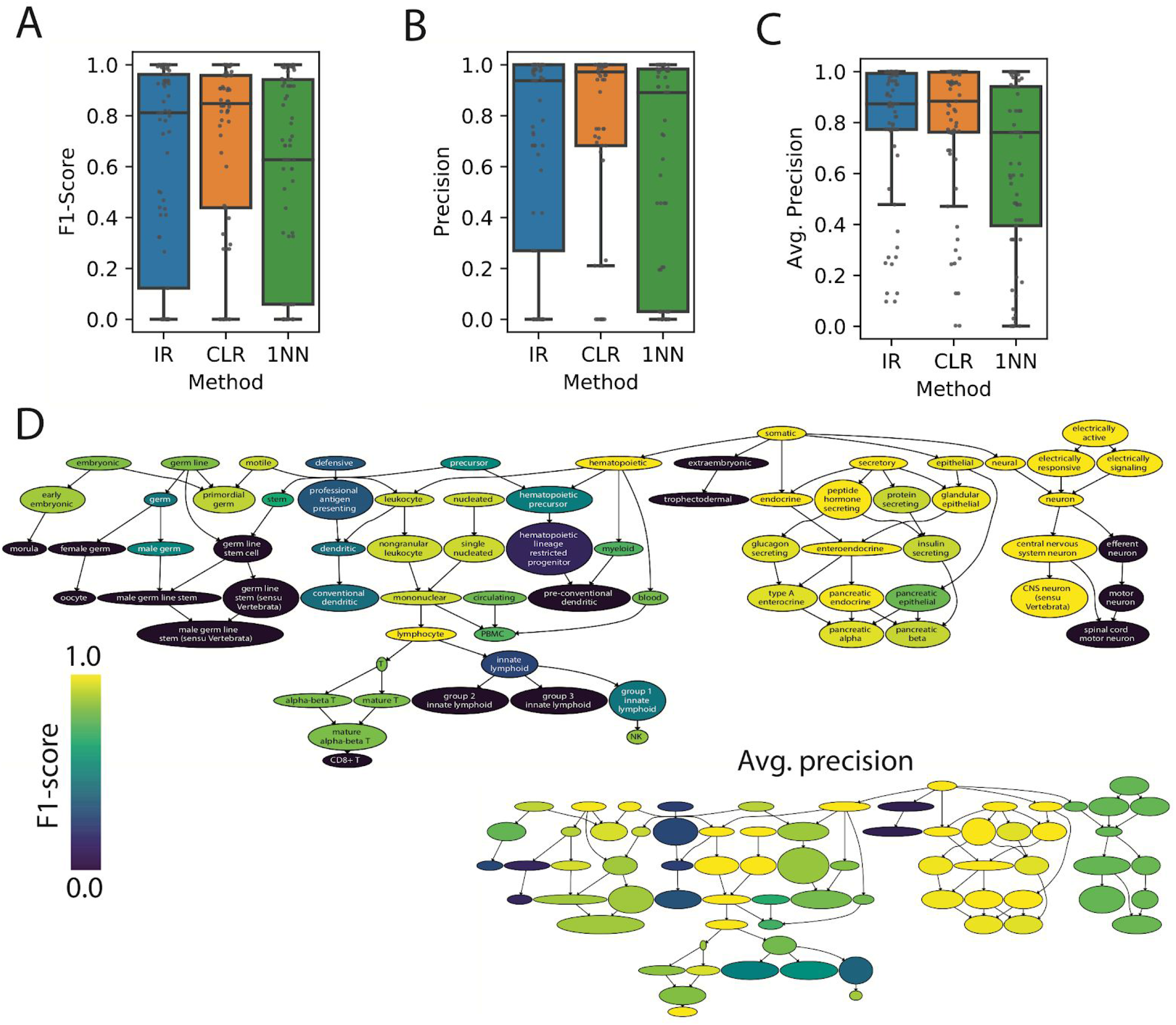
Results on non-droplet-based single-cell data. CellO’s performance on the 4,936 non-droplet-based cells considering only cells whose cell types are present in the bulk RNA-seq training set. We compare the distributions of (**A**) F1-score (**B**) precision, and (**C**) average-precision across all such cell types. (**D**) The subgraph spanning the non-droplet-based cells where each cell type is colored according to CellO’s (IR) F1-score (top) as well as by average-precision (bottom).

We also compared CellO to two existing methods, scMatch (Hou et al., 2019) and SingleR (Aran et al., 2019). scMatch and SingleR are most comparable to CellO because they come packaged with comprehensive reference datasets of human cells. Like CellO, these methods are designed to run out-of-the-box on diverse single-cell datasets. scMatch comes packaged with a reference dataset comprising data from the FANTOM5 project (Lizio et al., 2017). SingleR comes packaged with two comprehensive human reference datasets: a dataset comprising data from the Blueprint (Fernández et al., 2016) and ENCODE (Sloan et al., 2016) projects, and a reference set from the Human Primary Cell Atlas (Mabbott et al., 2013). To enable a comparison between scMatch, SingleR, and CellO, we project the outputs of scMatch and SingleR onto the Cell Ontology in order to evaluate scMatch and SingleR within the hierarchical classification framework. Specifically, for a given cell annotated by one of these methods with some cell type *C*, we also annotate the cell with all ancestors of *C* according to the Cell Ontology. First, we note that CellO’s training set included over 50% more of the cell types in this test set compared to those of scMatch and SingleR. In fact, most cell types in this test set, such as pancreatic islet cell types, are absent from scMatch and SingleR’s reference sets, thus indicating that a user would be required to supply their own reference set for annotating these cell types (**Figure 4A**). We thus evaluated each method on only cell types that exist in each respective method’s prepackaged reference set and found that CellO outperformed existing approaches (**Figure 4B**). We note that in this analysis, for a given method, we may remove from the analysis a specific cell type that is absent from a given method’s training set (e.g. stomach epithelial cell), but we would keep an ancestral cell type term (e.g. epithelial cell) if the method’s training set contains a sample labelled with this ancestral term (e.g. a sample labeled as intestinal epithelial cell).

**Figure 4.**
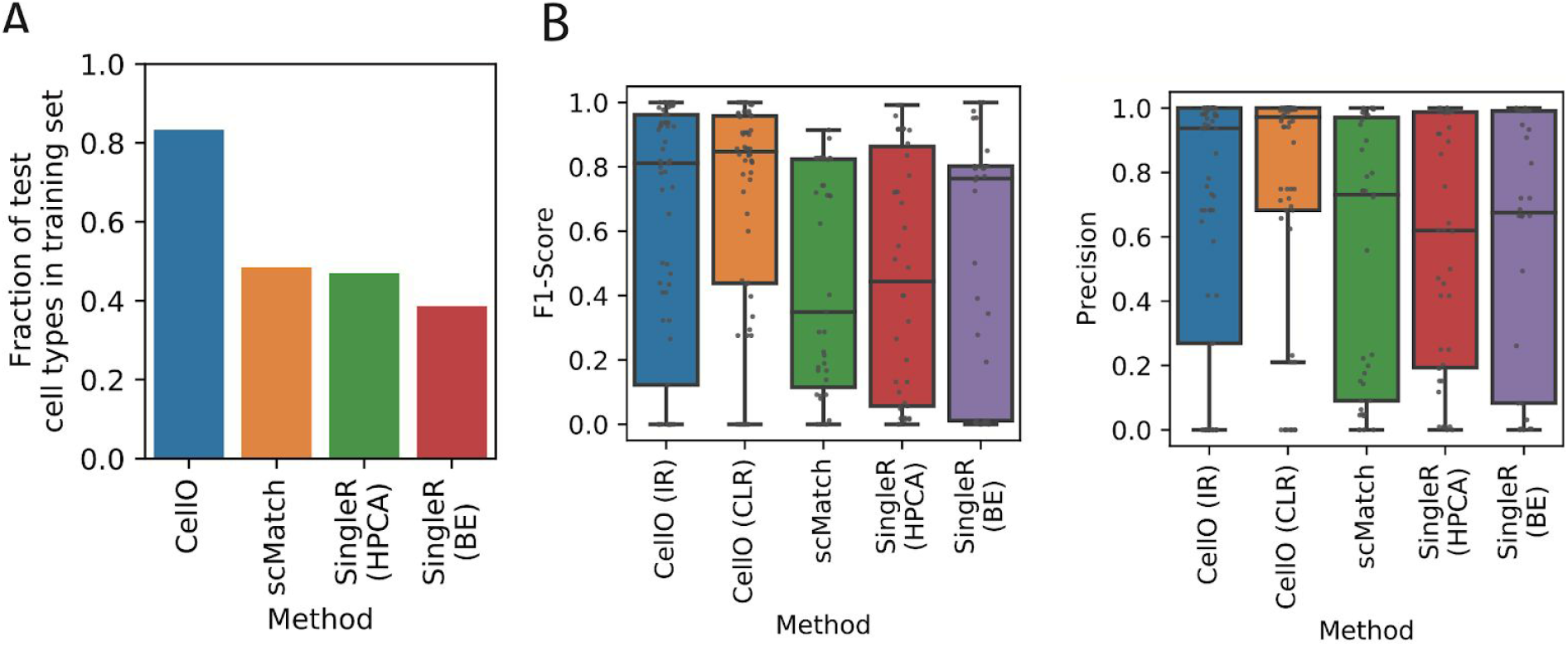
Comparison of CellO to existing approaches on non-droplet-based single-cell data. Evaluating CellO, SingleR, and scMatch on the non-droplet-based cells. (A) The fraction of cell types in the single-cell test dataset that are also present in each method’s training set. (B) The distribution of both F1-scores (left) and precisions (right) for only those cell types that are in each method’s training set. Note, each distribution evaluates different sets of cell types depending on the particular subset of cell types present in each method’s training set.

We further examined CellO’s classifications on two challenging subsets of the 7,366 cells that contained cells for which their combination of cell types was absent from CellO’s training set. First, we examined CellO’s accuracy on 1,978 healthy pancreatic islet cells from (Segerstolpe et al., 2016), which includes cell types that are absent from the bulk RNA-seq training set, specifically, ductal cells, acinar cells, epsilon cells, and delta cells. We found that CellO was able to correctly annotate the acinar cells as glandular epithelial cells, which is an ancestral cell type to acinar cells in the Cell Ontology (**Figure 5A, S3B**). This highlights the advantage of classifying against a hierarchy in that it enables CellO to annotate cells with a term higher in the ontology DAG when it is unsure about a cell’s more-specific cell type. We also note that a number of cells were uncharacterized in the original study due to not meeting a stringent quality control filter. CellO annotated many of these cells as pancreatic A cells (a.k.a. pancreatic alpha cells), which is plausible owing to both their close position to annotated A cells according to UMAP, which is known to preserve some level of global structure in high dimensional data (Becht et al., 2018), as well as the fact that A cells were found to be the most abundant endocrine cell type in Segerstolpe et al. (2016) of those that passed their stringent quality control filtering.

**Figure 5.**
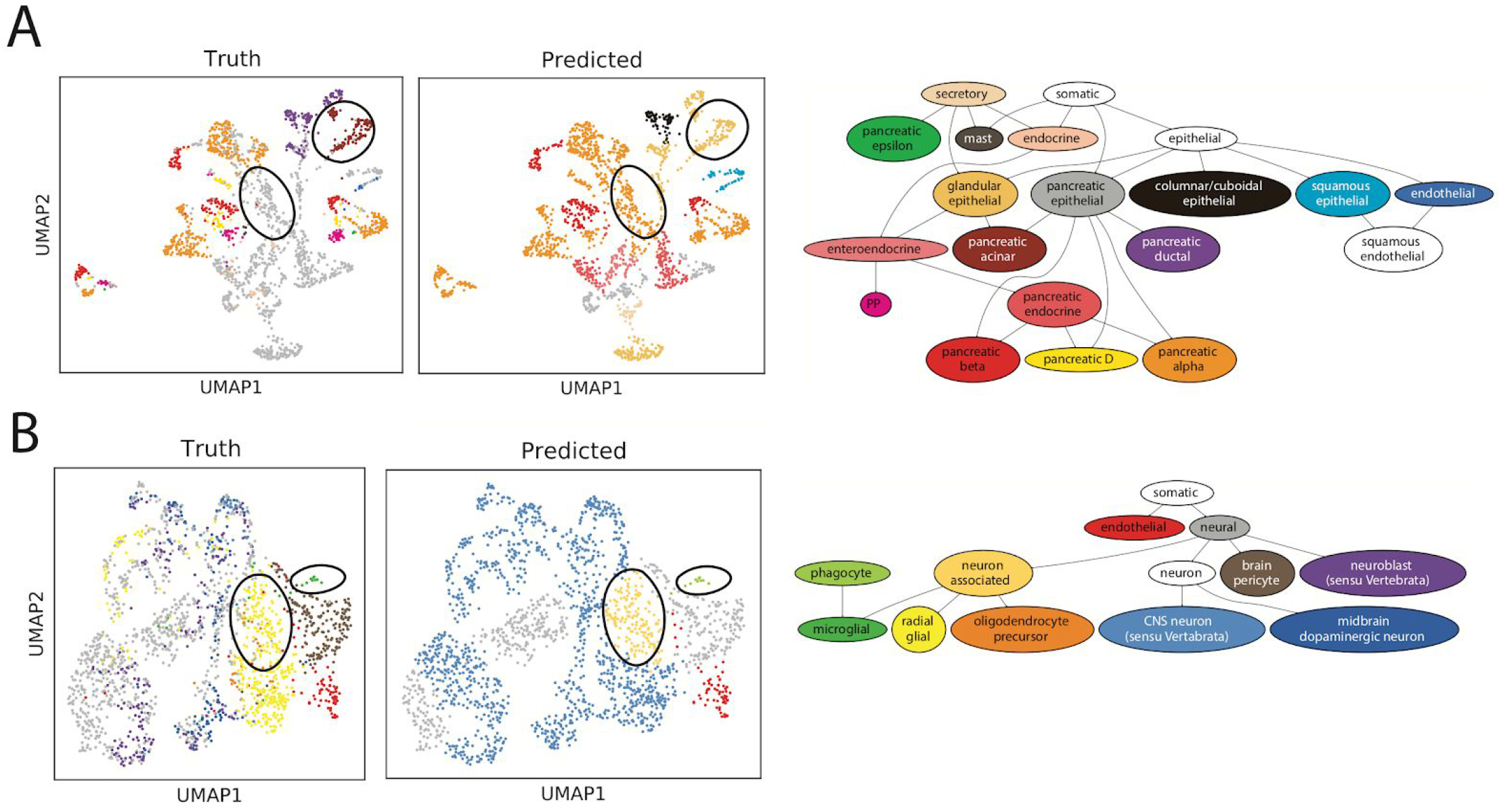
Examination of CellO’s performance on difficult datasets. (**A**) UMAP plots of all healthy cells in Segerstolpe et al. (2016) including cells for which their specific cell types are not present in Cello’s bulk RNA-seq training set. Cells are colored according to their true cell type (left) and (IR) predicted cell type (right). Highlighted are CellO’s predictions made on pancreatic acinar cells (top-right ovals) as well as a subset of uncharacterized pancreatic epithelial cells predicted as A cells (center ovals). (**B**) UMAP plots of human, embryonic neural cells from La Manno et al. (2016). Cells are colored according to their true cell type (left) and predicted cell type (right). Highlighted are CellO’s predictions made on both the microglial and glial cells and note that CellO annotates these cells using terms that are higher in the ontology’s graph than their true terms.

We also further examined CellO’s classification on 1,977 fetal neural cells from La Manno et al. (2016). Although the bulk RNA-seq training data contains samples of both embryonic cells and cells of various neural cell types, it does not contain any sample labeled as *both* neural cell *and* embryonic cell. Despite this discrepancy, CellO was able to annotate these cells with reasonable cell type labels (**Figure 5B, S2C**). We note that the microglial cells were annotated as phagocytes, which are an ancestral term to microglial cells in the Cell Ontology. Similarly, CellO annotated the glial cells as neuron associated cells. These examples again highlight CellO’s ability to annotate cells with a term higher in the ontology DAG when it is unsure about a cell’s more-specific cell type.

Finally, we evaluated CellO on FAC-sorted peripheral blood mononuclear cells from (Zheng et al., 2017) that were sequenced with Chromium 10x. We selected this dataset because it is one of the few droplet-based datasets for which the cell type labels are determined phenotypically (via sorting) rather than computationally (via expression analysis). We subsampled 2,000 cells from each of the ten sorted cell types and aggregated these cells together creating a dataset consisting of 20,000 cells. We first compared IR and CLR to the 1NN baseline and again found that IR and CLR outperformed 1NN with respect to median F1-score, although the difference between IR/CLR and 1NN was smaller than in the comparison on the aforementioned non-droplet-based scRNA-seq dataset (**Figure 5D**). Specifically, IR, CLR, and 1NN produced median F1-scores of 0.97, 0.96, and 0.95 respectively across all cell types.

Next, we compared CellO to scMatch and SingleR. In this analysis, we also ran SingleR using an immune-specific reference set of purified immune cells from Monaco et al. (2019). The cell types in this dataset are better represented in scMatch and SingleR’s respective reference sets, and thus, a comparison between CellO and these methods on this data better isolates performance differences between these methods. Like scMatch and SingleR, CellO struggled to accurately classify the T cell subtypes (**Figure 6A-C, S2C, S3**). Among the methods compared here, SingleR with the Monaco et al. reference set most accurately classified the T cell subtypes (**Figure S3**), though we note that this reference set is specialized for immune cell types whereas the other reference sets, including CellO’s, are more broad. We also note that CellO produced high average precision scores on most cell types including many of these T cell subtypes. Again, this indicates that CellO’s classifiers have learned to discriminate between these cell types; however, the threshold for calling these cell types may be non-optimal.

**Figure 6.**
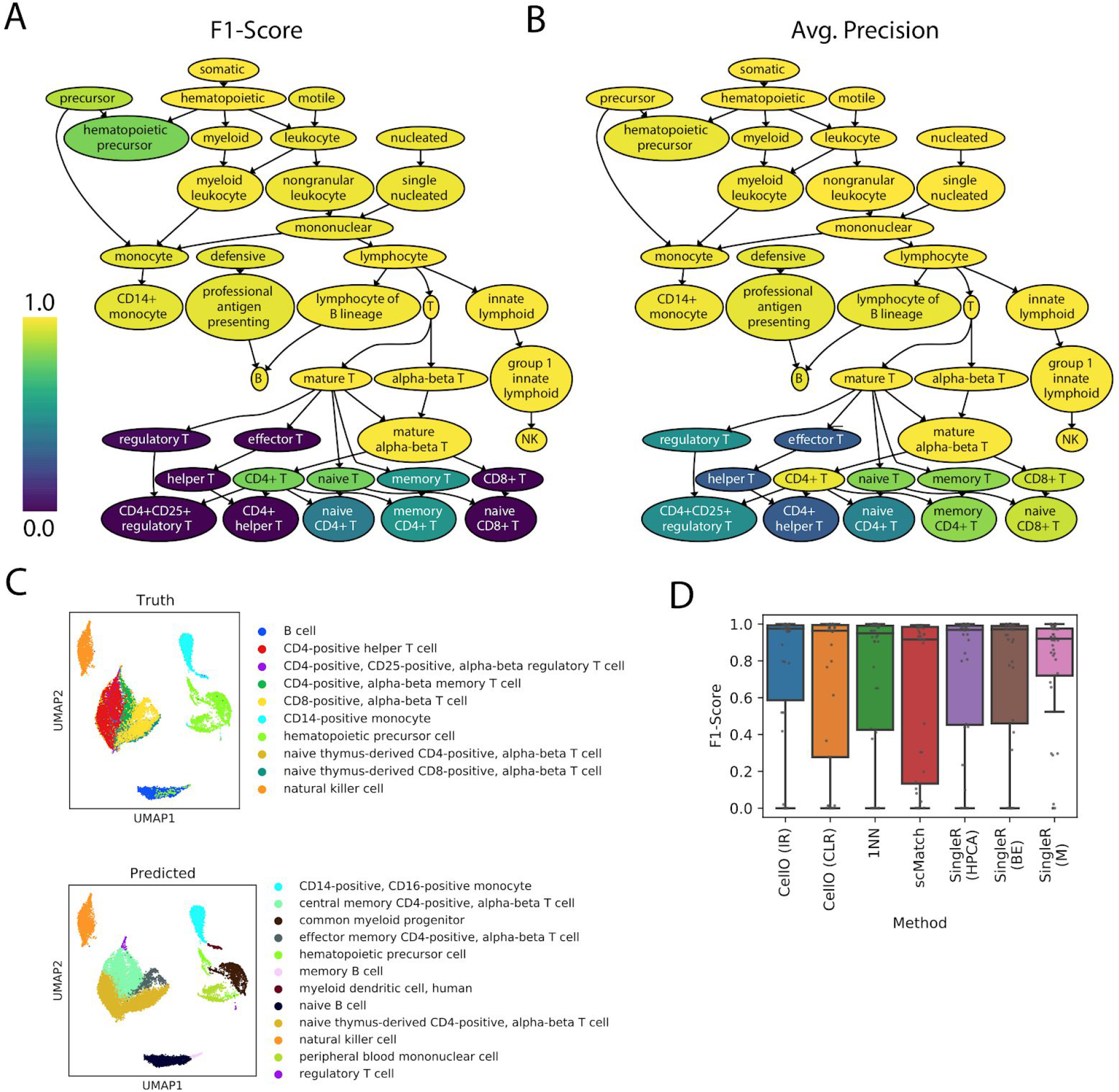
Results on 10x PBMC data. (**A**) The subgraph of the Cell Ontology spanning the 10x PBMC dataset from Zheng et al. (2017). Each cell type is colored according to CellO’s (IR) F1-score as well as (**B**) average-precision. (**C**) UMAP plots of the single-cell dataset where cells are colored by their true cell type (top) as well as the most-specific predicted cell type (i.e. lowest in the ontology) as output by CellO (bottom). (**D**) Boxplots displaying the distribution of F1-scores across all cell types for IR, CLR, 1NN, SingleR with the Human Primary Cell Atlas (HPCA), SingleR with the Blueprint+ENCODE reference (BE), SingleR with the Monaco et al. reference (M), and scMatch.

### User-friendly software

We provide a Python package for running CellO, using either IR or CLR, on a user-provided gene expression matrix (https://github.com/deweylab/CellO). CellO reduces the burden of reformatting and preprocessing an input expression matrix by accepting a variety of input file formats, including comma or tab-separated text files and HDF5, and by accepting expression data in a variety of units including counts or transcripts per million (TPM). To address the scenario in which the input dataset’s genes do not match those expected by the pre-trained classifiers, we provide functionality for a user to re-train the models on the bulk RNA-seq training set with a custom gene set. On a personal laptop, training a new classifier took 31 minutes to train IR and 11 minutes to train CLR on the full set of 58,243 GENCODE genes. Training time is reduced when trained on a smaller set of genes (e.g. only protein-coding genes). Finally, we note that relative performance of IR and CLR varies across cell types. To guide a user on their selection of either IR or CLR, we provide the average precision values achieved by both methods on each cell type in the bulk RNA-seq validation set (**Table S1**). We also provide average precision values and F1-scores achieved by both methods on each cell type on the test set of 4,936 non-droplet-based single-cells whose cell types were present in the bulk RNA-seq training set (**Table S2**).

### Interpretability of models

CellO makes extensive use of linear models, which are particularly amenable to interpretation especially when the coefficients are sparse (Gleicher, 2013). Although CellO’s models are not regularized to be sparse (as in Gleicher 2013), we sparsify them by selecting the top-ten genes per cell type according to the magnitude of the coefficients associated with each gene within each cell type’s one-vs.-rest binary classification model, which is used for CellO’s IR classifier. To enable their interpretation, we present a web-based tool, the CellO Viewer, for exploring these discriminative genes uncovered by the models (https://uwgraphics.github.io/CellOViewer/). The tool supports two modes of operation: a *cell-centric* mode (**Figure 7A**) and a *gene-centric* mode (**Figure 7B**). In the cell-centric mode the user can select cell types via a graphical display of the Cell Ontology in order to view and compare the most important genes for distinguishing those cell types. In the gene-centric view, the user can select genes and explore which cell types these genes are most important for distinguishing from the remaining cell types. The CellO Viewer uses an interactive display of the Cell Ontology’s graph to enable the user to navigate between cell types across the ontology.

**Figure 7.**
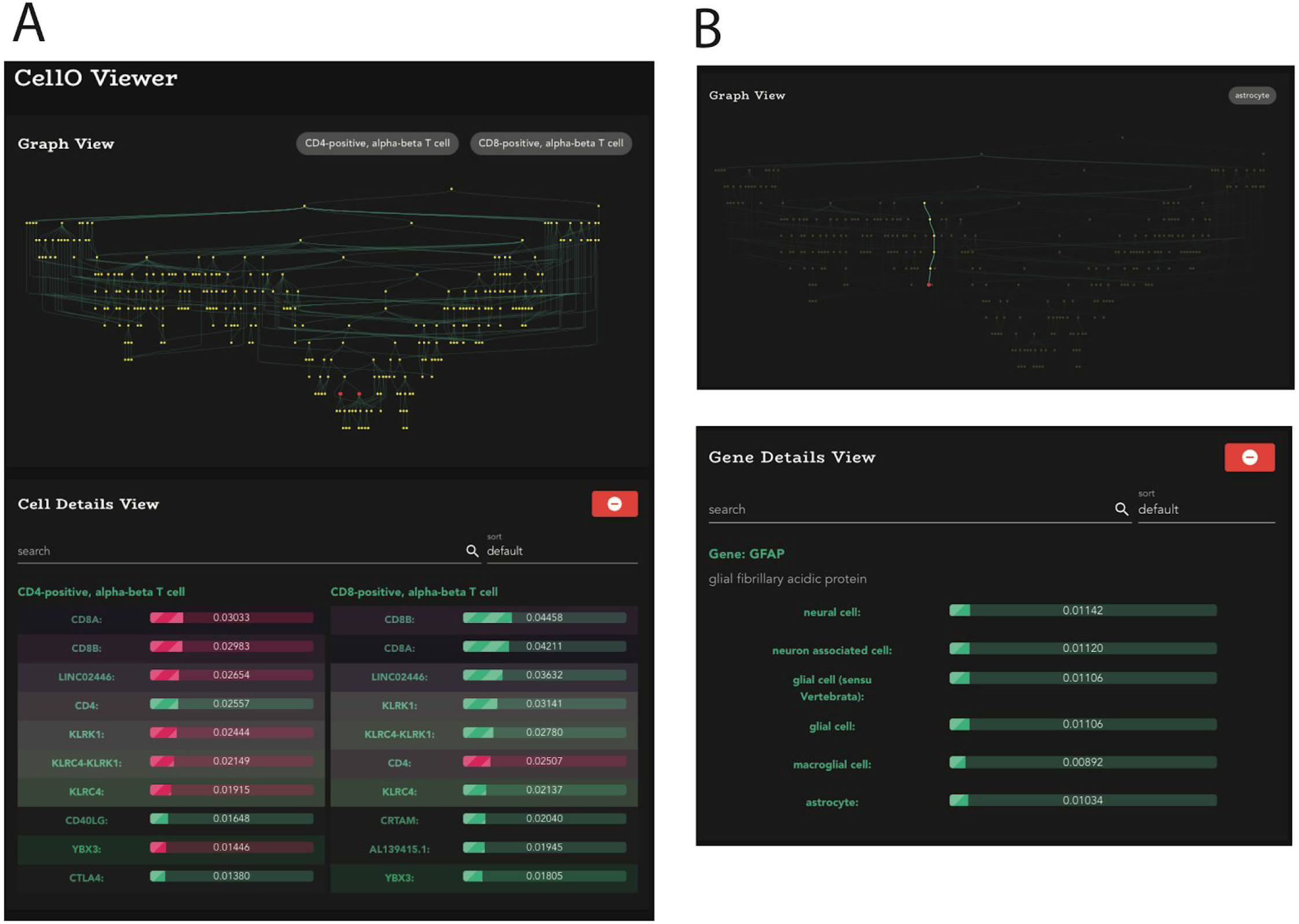
The CellO Viewer. Screenshots of the CellO Viewer web application for enabling the exploration of cell type-specific expression signatures across the Cell Ontology. (**A**) Comparing the top-ten genes between CD4+ T cells and CD8+ T cells (red nodes in the Graph View) ranked by the magnitude of their coefficients in their corresponding models. Genes that are shared between the two lists are highlighted with the same color. The CellO Viewer displays genes whose expressions are both positively correlated (green) and negatively correlated (red) with the selected cell types. (**B**) A screenshot of the gene-centric mode of the CellO Viewer with GFAP, an astrocyte marker, selected. For a given selected gene, the CellO Viewer will display the cell types within the DAG (top) and in list-form (bottom) for which the selected gene appears within the top-ten genes ranked by each model’s coefficients.

We found that across diverse cell types, many known cell type-specific marker genes were recovered by the CellO models and are presented by the CellO Viewer. For example, CD3D, CD3E, and CD3G, which are canonical markers for T cells, were all present within the top ten genes ranked according to the magnitude of their coefficients within the logistic binary regression model used for distinguishing T cells from all other cell types. Similarly, CD4 and CD8 were present in the top genes for the CD4+ T cell and CD8+ T cell models, respectively (**Figure 7A**). In a more complex example, the genes GCG, LOXL4, DPP4, GC, and FAP, known markers for pancreatic alpha cells, and INS, IAPP, and ADCYAP1, known markers for pancreatic beta cells (Segerstolpe et al., 2016), all appear within the top ten genes for their respective cell types.

Interestingly, certain genes appear in the top ten coefficients for broad cell types, but not more specific cell types, indicating that CellO is able to find signals specific to broad cell type categories. For example, DDX4 appeared in the top ten genes for distinguishing germ line cells, but did not appear within the top ten genes for any of the more specific germ cell subtypes.

DDX4 is known to be expressed in germ cells across both sexes (Hickford et al., 2011). Similarly, the gene NRG1 appeared in the top ten genes for distinguishing precursor cells, but did not appear within the top ten genes for any of the more specific cell types that are descendents of precursor cells within the ontology. NRG1 is known to play a role in the development of a number of organ systems (Lemmens et al., 2007; Mei and Xiong, 2008).

## Discussion

In this work, we explore the application of hierarchical classification algorithms to cell type classification with the Cell Ontology using a novel, well-curated set of human primary cell RNA-seq samples. This dataset may prove useful for future investigations of cell type expression patterns or for use in cell type deconvolution methods (Aran et al., 2019; Newman et al., 2015). We demonstrate that the trained classifiers perform well across cell types in diverse single-cell datasets and outperformed existing cell type annotation methods when trained on their comprehensive reference sets. We packaged these classifiers into an easy-to-run Python package called CellO.

In our exploration of methods for correcting the independent one-vs.-rest classifiers, we found that discriminative methods outperformed the generative BNC approach implemented in URSA (Lee et al. 2013). We hypothesize that BNC suffers in comparison due to two causes. First, BNC’s probabilistic model makes strong assumptions regarding the generative process of classifier scores and true cell type assignments. Second, BNC requires estimating the conditional probability distribution of each classifier’s output scores (i.e. distance from the decision boundary) conditioned on the true cell type labels, which may be difficult to estimate accurately given the limited quantity of training data available for each cell type.

By using linear models, CellO’s trained parameters are easily interpreted as cell type specific signatures across the ontology. However, we note that since certain cell types undergo similar sorting and preparation procedures (e.g., fluorescence activated cell sorting), it remains unclear to what extent these procedures affect gene expression and thus confound cell type. We sought to mitigate this effect by using data from a diversity of studies. We also note that the CLR algorithm may help to further mitigate this effect, since the binary classifiers trained in this framework for each cell type condition on the sample belonging to the parent cell types. Thus, for a given cell type, if samples of its parent cell types were prepared through similar procedures, the learned model parameters for that cell type will better capture biological cell type signatures.

There are a number of avenues that require further investigation. First, we note that calibrating discriminative models trained on bulk RNA-seq data and applying them to single-cell data is challenging. In this work, we developed techniques for addressing this issue, but there remains a gap between the performance of CellO when evaluated with average precision versus when evaluated with F1-score. The very high average-precision scores across many cell types indicates that with better calibration, CellO’s accuracy when making binary yes-no decisions for each cell type could be improved. Future work will investigate alternative approaches to calibrating CellO’s models in order to improve CellO’s binary cell type decisions. Second, CellO was trained on only human samples. Future work will entail curating comprehensive training sets from the SRA for other species such as mouse.

Finally, we expect the performance of hierarchical classifiers to improve as both more data is collected and as the Cell Ontology is expanded. Most importantly, we expect the calibration of the classifiers to improve as more training data becomes available for each cell type. More data will be collected both as data is continually added to the SRA and as improvements are made to the SRA’s metadata thereby allowing retrieval of previously undiscovered primary cell samples.

## Acknowledgements

M.N. Bernstein thanks Gary H. Bernstein, Ron Stewart, and Christina Kendziorski for helpful conversations.

## Author Contributions

M.N.B. led the conceptual development of the ideas present in this article and implemented both the CellO software and experiments. C.N.D. supervised the conceptual development and implementation of the software and experiments. M.N.B. and C.N.D. wrote the manuscript. Z.M. implemented the CellO Viewer. M.G. supervised the development of the CellO Viewer. M.N.B., Z.M., M.G., and C.N.D. contributed to the conceptual design of the CellO Viewer.

## STAR Methods

### Methods Details

#### Data curation

To create the training set of primary cell bulk RNA-seq samples, we first selected all samples labelled as a “primary cell” sample by the MetaSRA (v1.4). Thus, we followed the conservative definition for a primary cell sample by Bernstein et al. (2017), which requires that a sample has not undergone passaging beyond the first culture. We followed this selection with a manual curation of each sample’s technical variables by consulting sources of metadata that are not captured by the MetaSRA annotation process, such as fields in Gene Expression Omnibus (Clough and Barrett, 2016) records and each study’s publication. In total, we annotated 27,097 samples (available to download at http://deweylab.biostat.wisc.edu/cell_type_classification/). We then removed all samples that were either incorrectly labelled as primary cell samples or had been experimentally treated. When found, we also corrected errors in the MetaSRA-provided Cell Ontology labels by both adding additional cell types that were missed by the MetaSRA as well as removing incorrect cell type labels. This curation effort resulted in the final set of 4,293 samples.

#### Data processing

We quantified the gene expression of all RNA-seq samples from the SRA (both bulk and non-droplet-based single-cell samples) with kallisto (v0.43.1) (Bray et al., 2016) using human genome release GRCh38 with GENCODE annotation version 27. We chose kallisto for gene expression quantification in order to prioritize processing speed on this large dataset, figuring that any small loss in accuracy (at the gene level) relative to a less approximate, but slower approach would not be significant for the cell type classification task. This produced estimated counts for 200,401 isoform-level genomic features. We summed the transcripts per million (TPM) values by gene to produce TPM’s for 58,243 gene-level features. The curated metadata and associated quantified samples are available to download at http://deweylab.biostat.wisc.edu/cell_type_classification.

#### Notation

In the following descriptions of the methods used in this work, we let ***x*** ∈ ℝ^*G*^ denote a gene expression profile, where *G* is the number of considered genes and ***x*** is measured in units of log(TPM+1) where TPM are transcripts per million. We let *n* denote the number of samples, *m* denote the number of considered cell types, *y_i,j_* ∈ {0, 1} denote the cell type assignment for cell type *j* ∈ [*m*] and sample *i* ∈ [*n*].

#### Binary classification with logistic regression

We use L2-penalized logistic regression for each binary classifier as implemented by scikit-learn (Pedregosa et al., 2011) using the LIBLINEAR solver (Fan et al., 2008). To speed up the training of each binary classifier, we preprocessed the bulk RNA-seq training data using principal components analysis (PCA) as implemented in scikit-learn. Specifically for each cell type *j*, each classifier is trained by minimizing the following loss-function:

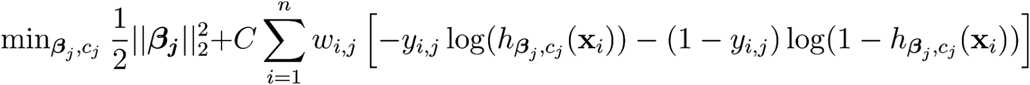

where

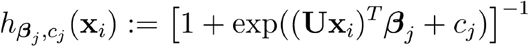

is the logistic function, ***β***_*j*_ ∈ ℝ^*k*^ are the model-coefficients for cell type *j*, *c_j_* is the intercept for cell type *j*, *C* controls the strength of the regularization, *n* is the number of training samples, *w_i,j_* is a per-sample weight for handling class-imbalance in cell type *j*’s model, and **U** ∈ ℝ^*k*×*m*^ is the PCA loadings matrix.

We also note that most training sets are highly unbalanced. We found that the models were better calibrated when the loss-function weighted each training sample such that the positive and negative samples contribute equally to the loss function (via the *w_i,j_* weights above) as evidenced by the improved F1-scores when using a threshold of 0.5 for making discrete yes-no cell type decisions (**Figure S5B**). We implemented this class-balancing by setting the *class_weight* parameter to ‘balanced’ in scikit-learn’s *LogisticRegression* class constructor.

#### Isotonic regression correction

We train a binary classifier for each cell type *j* ∈ [*m*] to model *p*(*y_j_* | ***x***) using logistic regression and a one-versus-rest training strategy. As proposed by Obozinski *et al* (2008), these probabilities are then reconciled with the ontology graph using isotonic regression. Specifically, we output the set of probabilities:

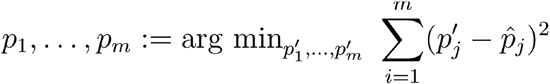

subject to

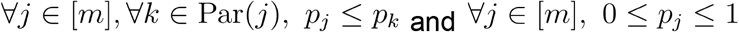

where ∀_*j*_ ∈ [*m*], 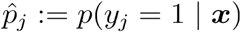 as output by each classifier and Par(*j*) is the set of parent cell types for cell type *j*. CellO uses the quadprog Python package (https://pypi.org/project/quadprog/) for solving this quadratic optimization problem.

#### Bayesian network correction

A binary classifier is trained for each cell type and a one-versus-rest training strategy. The classifier outputs are then reconciled with the ontology graph using a Bayesian network as proposed by Lee et al. (2013). The true assignments for each cell type within a given sample, denoted *y*_1_,…,*y_m_*, are modelled as latent random variables, and the classifier outputs, denoted *f*_1_(***x***),…,*f_m_*(***x***) (signed distances to each decision boundary), are modelled as observed random variables in a Bayesian network. The final output probability for cell type *j* is then the marginal probability

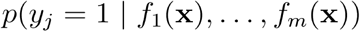

More specifically, for a given cell type *j*, we model the conditional distribution of the classifier’s output *f_j_*(x) (distance to the decision boundary) conditional on its true label *y_j_* as a discrete random variable constructed as follows. We partition the training data for cell type *j* into two folds ensuring that no study is split between folds while attempting to keep the sizes of the two folds as similar as possible. We then train on one fold and compute the classifier scores from the second fold (for each of the two folds).

Using all of these scores, we then compute a histogram where the bin sizes are determined using a second 2-fold cross-validation scheme. Specifically, we then test a number of bin sizes by first estimating a histogram density function using data in one fold and then computing the likelihood of the data in the second fold (performing this procedure for both folds). The histogram density function is given by

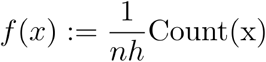

where *n* is the total number of data points, *h* is the width of each bin, and *Count*(*x*) is the number of data points sharing the same bin as *x*. We choose a bin size that maximizes the mean of the two data log likelihoods computed on each fold.

As described by Lee et al. (2013), the true cell type assignments *p*(*y*_1_,…,*y_m_*) factor according to the ontology graph:

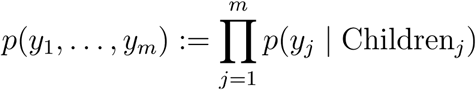

where

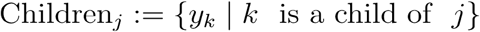

are the assignments to the children cell types of cell type *j* in the ontology. These conditional distributions *p*(*y_j_*|Children_*j*_) enforce consistency with the ontology by defining

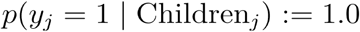

if 1 ∈ Children_*j*_. Otherwise,

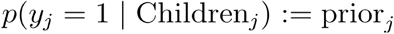

The prior_*j*_ values are computed from counts in the training data. Specifically, the prior for each leaf-node cell type *c* is simply the fraction of samples in the data set labelled as *c*. For each internal node *c*, the prior is computed as the fraction of all samples labelled as *c*, but not labelled as any child of *c*. A pseudocount of one was used in the calculation for all priors. Due to the size of the ontology, we perform approximate inference using Gibbs sampling rather than exact inference using the Laurintzen algorithm as was performed by Lee et al.

To test the impact of the graph-structured prior on this algorithm, we also tested a naive Bayes variant in which each cell type is predicted independently (i.e., without the graph-structured prior). We found a significant improvement in performance for the graph-structured BNC algorithm over the naive Bayes algorithm indicating that the graph structured prior is an important component of this algorithm (**Figure S6**).

#### True Path Rule

We train a binary classifier for each cell type *j* ∈ [*m*] to model *p*(*y_m_* | ***x***) using logistic regression and a one-versus-rest training strategy. As proposed by Notaro et al. (2017), this method involves two passes across the ontology: on a bottom-up pass, each cell type’s output probability is averaged with the outputs of all child cell types classifiers for which the classifier makes a positive prediction according to a predefined threshold. More specifically, each cell type *j*’s output probability is set to

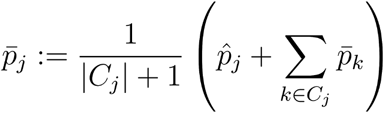

where 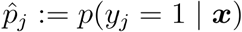 according to the classifier and

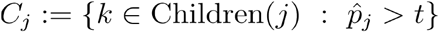

is the set of children of cell type *j* for which the classifier output a positive prediction according to a predefined threshold *t*. We used a threshold of *t* = 0.5. This bottom-up pass allows for sharing of information across the classifiers. In the top-down pass of the ontology, the output probabilities are set to ensure consistency with the ontology. This procedure works as follows: each node in the ontology is visited according to the topologically sorted order of nodes and for a given visited cell type *j*, its final probability is set to

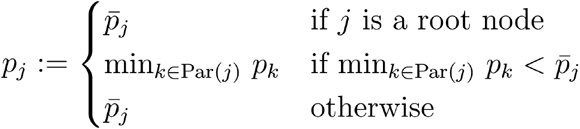

where *Par*(*j*) are the parent nodes of node *j* in the DAG.

#### Cascaded logistic regression

Classification is made in a top-down fashion starting from the root of the ontology downward as proposed by Obozinski et al. (2008). This is accomplished by training a logistic regression, binary classifier (although any binary classifier that outputs a prediction probability can be used) for each cell type *j* ∈ [*m*] to model the distribution

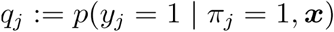

where *π_j_* ∈ {0, 1} indicates whether the sample belongs to all of the parents of *j* in the ontology. In order to model these distributions, each cell type’s negative training examples consist of those samples that are labeled with all parent cell types, but not the target cell type.

Given these learned distributions, the probability that ***x*** originates from cell type *j* is computed via

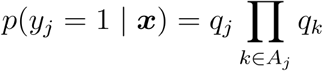

where *A_j_* denotes the ancestors of cell type *j* in the ontology’s DAG.

#### One-nearest neighbors

Given a query gene expression profile ***x***, we return all cell type labels belonging to the training set expression profile

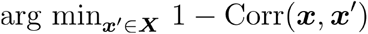

where Corr(***x***, ***x***′) is the Pearson correlation of the expression values in ***x*** and ***x***′ as implemented in Python’s SciPy package (Virtanen et al., 2020).

### Quantification and statistical analysis

#### Partitioning bulk RNA-seq data into training and test sets

In order to find the optimal parameters and configurations for CellO, we partitioned the bulk RNA-seq training dataset into a pre-training set and validation set (**Figure S4**). When creating this partition, we sought to satisfy a number of criteria that would enable unbiased estimation of performance across cell types. First, we required that no study be split between the pre-training and validation sets in order to ensure that a model is never tested on data from a study on which it was trained. This mitigates the possibility that the algorithm will provide an overly optimistic estimate of the generalization error when run on the validation set. Second, we sought an approximately 80/20 split of the data between the pre-training and validation sets. Third and finally, we sought for all cell types to be represented in both the pre-training and validation sets. We framed this partitioning task as an optimization problem where our four criteria were encoded in an objective function. Minimizing this objective function entails creating a partition that most closely meets the aforementioned four criteria. To optimize the objective function, we performed a simple hill-climbing procedure where we moved a study’s data from the validation set to the pre-training set if such a move resulted in a partition that improved the objective function.

#### Parameter tuning

To choose the optimal number of principal components, we evaluated various numbers of principal components by training the collection of one-vs.-rest classifiers on the bulk RNA-seq pre-training set and evaluating on the bulk RNA-seq validation set. We found that using 3,000 principal components performed equally well to using the raw gene expression values as features. Further, we found performance degraded as the number of principal components decreased (**Figure S5A**). We then performed a parameter sweep of penalty-weight parameters (i.e. regularization-strength parameters) for IR and CLR and chose the values that maximized the median average precisions for each method (**Figure S5D**).

#### Model calibration

We found that for many cell types, there were relatively few studies in the training set for which samples of these cell types originated and thus, we found that the models tended to overfit to the training set leading to poorly calibrated models. This was evidenced by the fact that the average-precision scores were high across most cell types; however, the F1-scores were much lower when using a default threshold of 0.5 for making binary cell type decisions (**Figure S5D**). The high average precision scores indicate that the positive and negative examples of each cell type are well separated in the ranking of samples when ranked according to the classifier output probability for the given cell type. However, the poor F1-scores indicate that the default threshold of 0.5 is non-optimal for separation.

To address this issue, we used a data-driven approach to find empirical thresholds according to a leave-study-out cross-validation experiment. Specifically, we performed leave-study-out cross-validation on the pre-training set of bulk RNA-seq samples and used these results to empirically choose a threshold for each cell type such that if a given threshold less than 0.5 led to a higher F1-score than the default threshold of 0.5, we select this empirical threshold. When applying these data-driven thresholds to the validation set, we observed a significant increase in the F1-scores across cell types under the independent one-vs.-rest classifiers approach (**Figure S5B**). Thus, when training the final CellO models (both CLR and IR), we performed this same cross-validation procedure on the entire bulk RNA-seq training set to select data-driven thresholds.

We note that even with these selected thresholds, CellO often output more than one specific cell type for a given sample (e.g. both natural killer cell and T cell), which may confuse a user of the tool. This phenomenon is a consequence of both mis-calibrated models (despite the data-driven thresholding procedure) and because CellO performs multi-label hierarchical classification. To address this issue, whenever CellO outputs more than one specific cell type, we select only the cell type with highest output probability along with all ancestor cell types. All F1-scores reported in the main text followed from this correction procedure and thus, are an apt measure of the practical utility of CellO on real data. This procedure is illustrated schematically in Figure S7.

#### Model interpretation

We note that although the logistic regression model’s coefficients ***β***_*j*_ ∈ ℝ^*k*^ (where *j* denotes the index for a given cell type) weight the principal components rather than genes, each gene’s contribution to the model’s decision can be recovered by

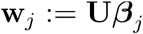

where **U** ∈ ℝ^*m*×*k*^ is the PCA loading matrix **w**_*j,g*_ and describes the contribution of gene *g* to the model’s decision. The full set of these vectors over all cell types can be explored within the CellO Viewer web application.

#### Robustness of CellO to single-cell clustering parameter

In order to test CellO’s robustness to cluster granularity when clustering single-cell data, we tested CellO on the Zheng et al. PBMC dataset using differing values for Leiden’s resolution parameter. We found both CLR and IR to perform similarly across various values of Leiden’s resolution parameter (**Figure S8**).

#### Evaluation metrics

For a given input dataset, CellO generates two sets of outputs: binary yes-no decisions for each cell type assignment as well as probability scores that quantify how likely each cell should be assigned to any given cell type. Both of these outputs can be represented as matrices. More specifically, given an input dataset, **X** ∈ ℝ^*n*×*G*^ where is the number of cells and *G* is the number of genes, the binary yes-no decisions can be represented as a matrix ***B*** ∈ {0, 1}^*n*×*m*^ where *m* is the number of cell types and *B_i,j_* = 1 if the classifier predicts cell to be of cell type *j*. The probability scores can also be represented as a matrix *S* ∈ [0, 1]^*n*×*m*^ where *S_i,j_* denotes the classifier’s confidence that cell type *i* should be labelled as cell type *j*. Finally, the true cell type assignments can also be represented as a matrix *T* ∈ {0, 1}^*n*×*m*^ where *T_i,j_* = 1 if cell *i* is truly of cell type *j*.

We first define metrics for comparing *B* to *T*. For cell type *j*, we define the number of true positives (TP), false positives (FP), and false negatives (FN) as:

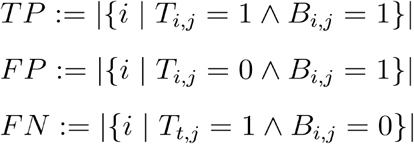

We then evaluate the classifier’s performance on cell type *j* using precision, recall, and F1-score, which are defined as

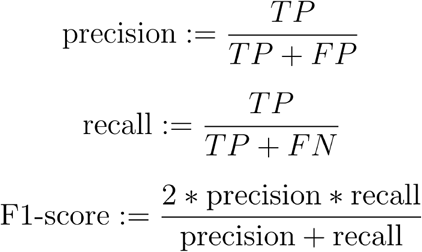

We also evaluate the methods using “joint” precision-recall curves in which each cell-cell type pair is considered to be an independent prediction (i.e. each element of *S*). These pairs are then ranked and the corresponding precision-recall curve is constructed (Obozinski et al. 2008).

We note that for a given cell, the ground-truth assignments for cell types that are more-specific than the cell’s most-specific ground-truth cell type are ambiguous since these samples could, in theory, truly be of a more specific cell type than they are labelled with (e.g., a cell labelled as a T cell could be a CD8+ T cell even if it isn’t annotated as such). Thus, when computing the aforementioned metrics for a given cell type within these single cell datasets, we exclude those cells that are labelled most-specifically as an ancestor of the cell type. For example, for metrics calculated for CD8+ T cells, we would exclude from the calculations those cells that are most-specifically labelled as T cells.

### Resource Availability

#### Lead Contact

Further information and requests for resources should be directed to and will be fulfilled by the Lead Contact, Colin Dewey.

#### Data and Code Availability

A Python package for running CellO can be found at https://github.com/deweylab/CellO. The data used in this work can be found at http://deweylab.biostat.wisc.edu/cell_type_classification/. The CellO Viewer can be accessed at https://uwgraphics.github.io/CellOViewer/. The code implementing the CellO Viewer can be found at https://github.com/uwgraphics/CellOViewer. All code for performing the experiments in this work can be found at https://github.com/deweylab/cell-type-classification-paper.

#### Materials Availability Statement Examples

This study did not generate new unique reagents.

---

**Supplementa1 Figure 1.**
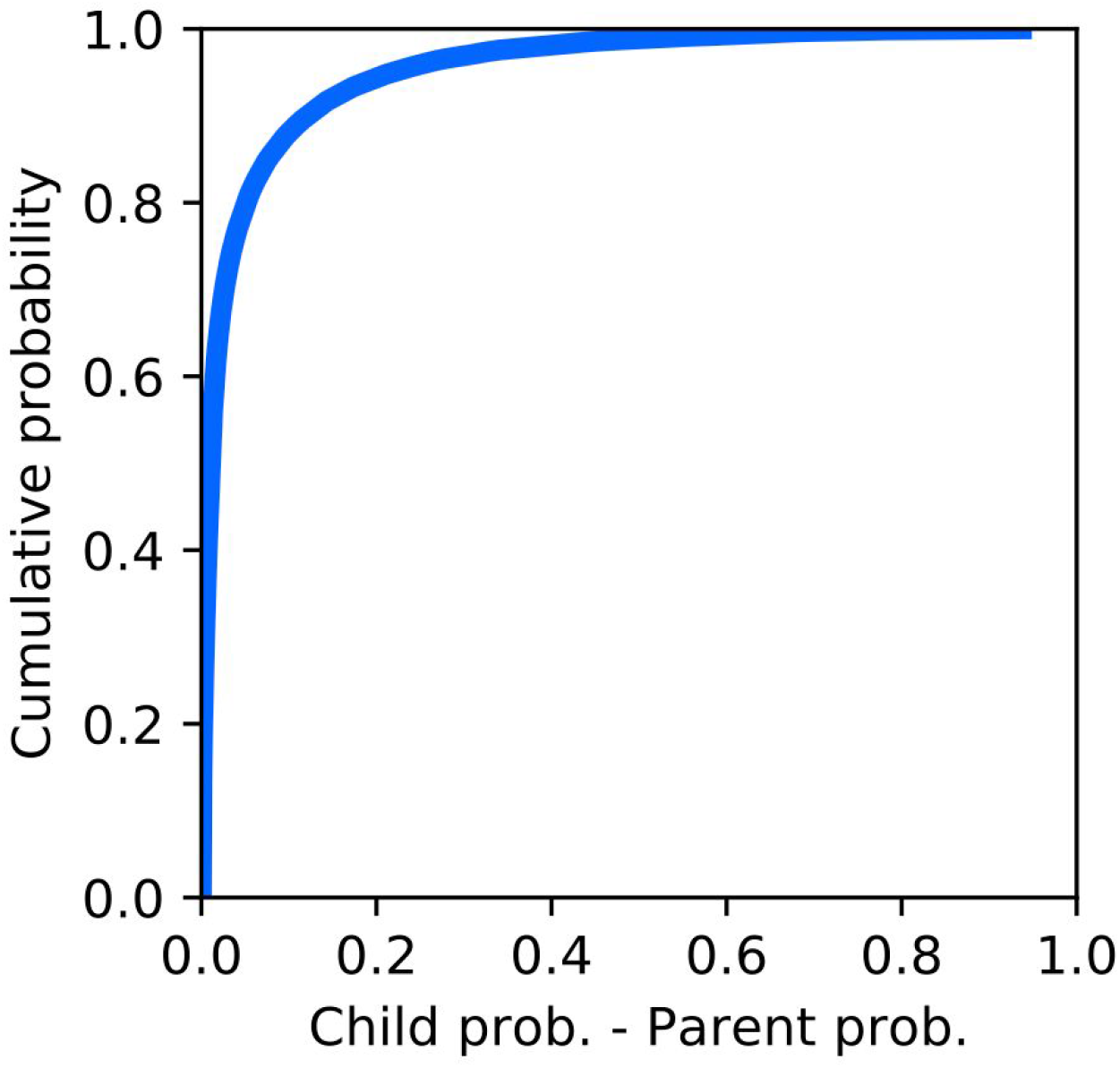
Distribution of edge inconsistencies. The cumulative distribution function over the difference in probability between the parent and child classifiers for all edges for which either the parent or child classifier output a probability greater than 0.01.

**Supplementa1 Figure 2.**
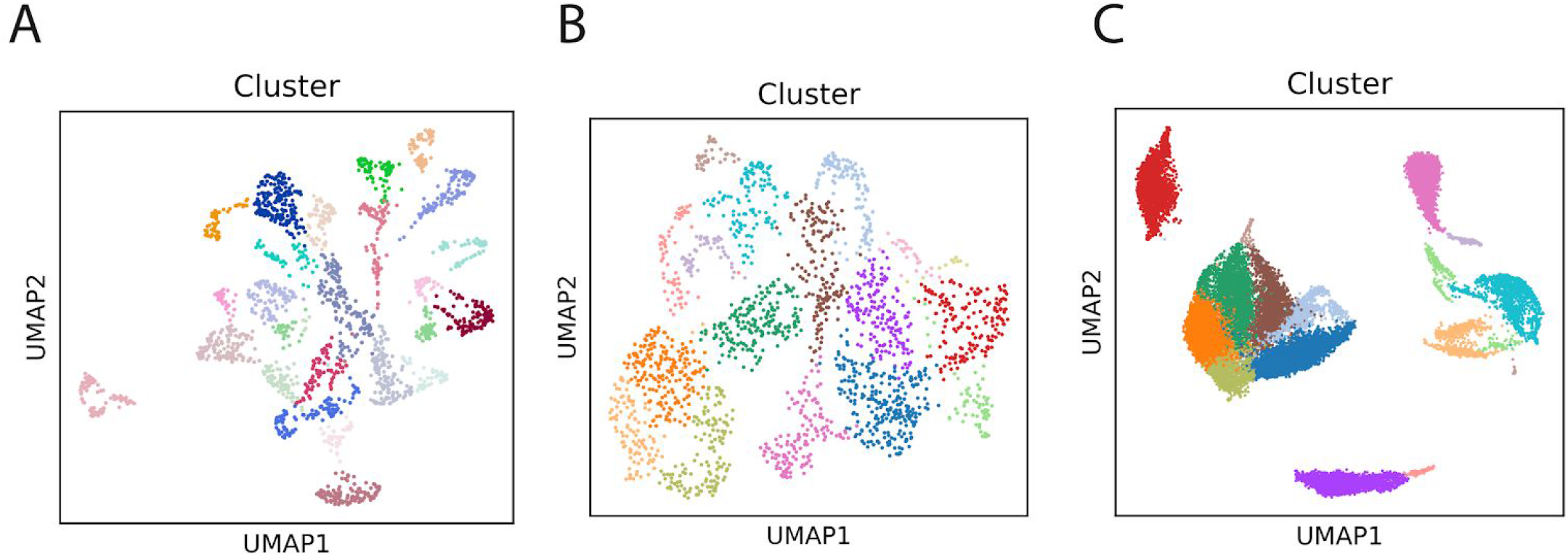
Cluster assignments for single-cell datasets evaluated in this work. UMAP plots of the single-cell datasets examined this work where each cell is colored according to its CellO cluster assignment. These clusters were aggregated to compute a mean expression profile and on which CellO was run. Plots are displayed for (**A**) the Segerstolpe et al. (2016) dataset of healthy pancreatic cells, (**B**) the La Manno et al. (2016) dataset of fetal neural cells, and (**C**) the Zheng et al. (2017) dataset of PBMCs.

**Supplementa1 Figure 3.**
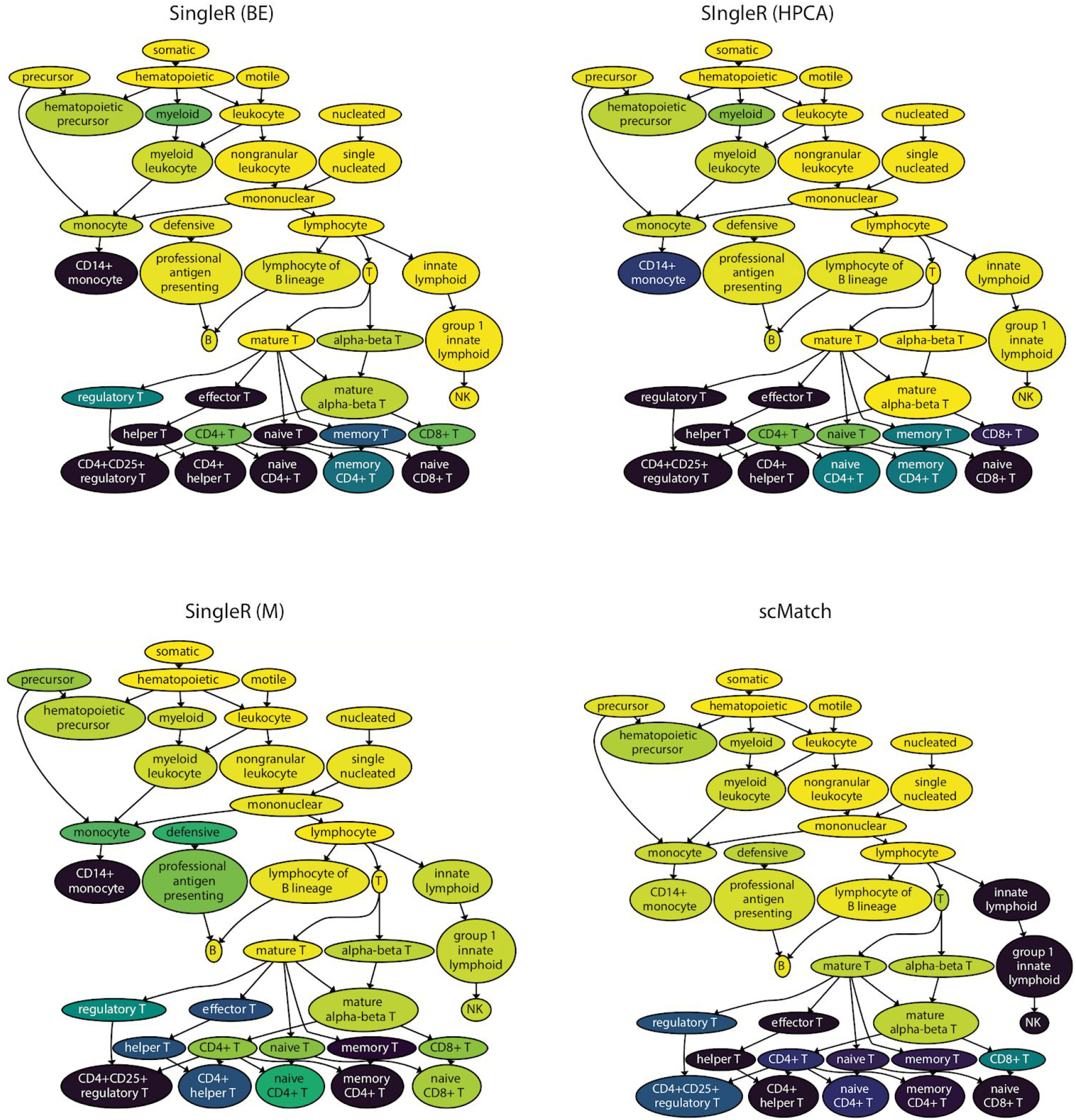
PBMC F1-scores produced by existing methods. The subgraph spanning the PBMC’s from Zheng et al. (2017) where each node is colored according to the F1-scores produced by SingleR with the Blueprint+Encode reference (BE), SingleR with the Human Primary Cell Atlas reference (HPCA), SingleR with the Monaco et al. (2019) reference (M), and scMatch.

**Supplementa1 Figure 4.**
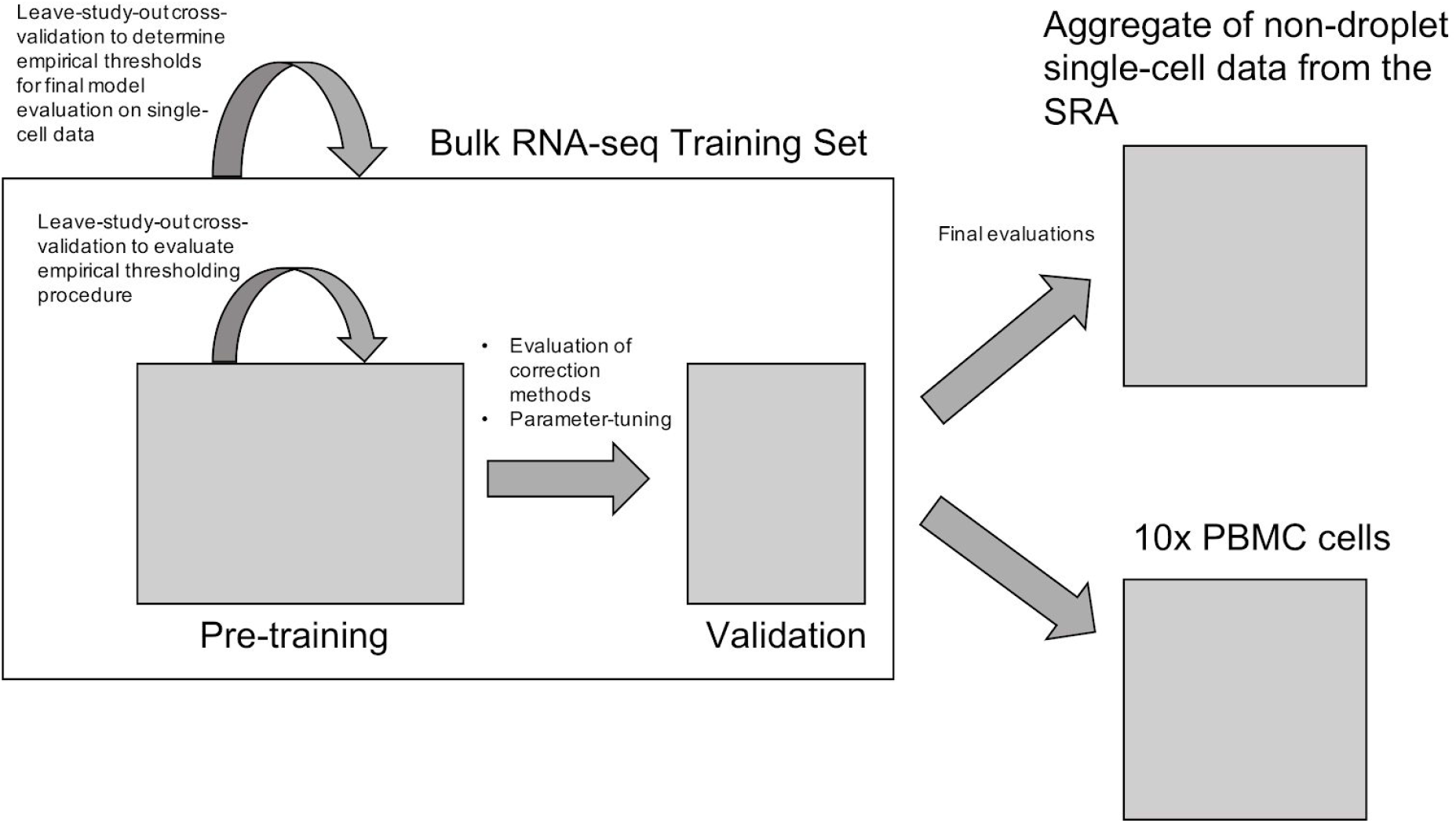
Overview of training and testing sets. A schematic illustration of the training and testing sets used in this study. A bulk RNA-seq dataset was split into a pre-training and validation set for tuning the parameters of the binary classifiers as well as for evaluating the graph-correction methods. The full bulk RNA-seq dataset was then used to train the final models that were then evaluated on two sets of scRNA-seq data. The first set consisted of an aggregation of diverse non-droplet-based datasets from the SRA. The second dataset consisted of FAC-sorted PBMCs from Zheng et al. (2017).

**Supplementa1 Figure 5.**
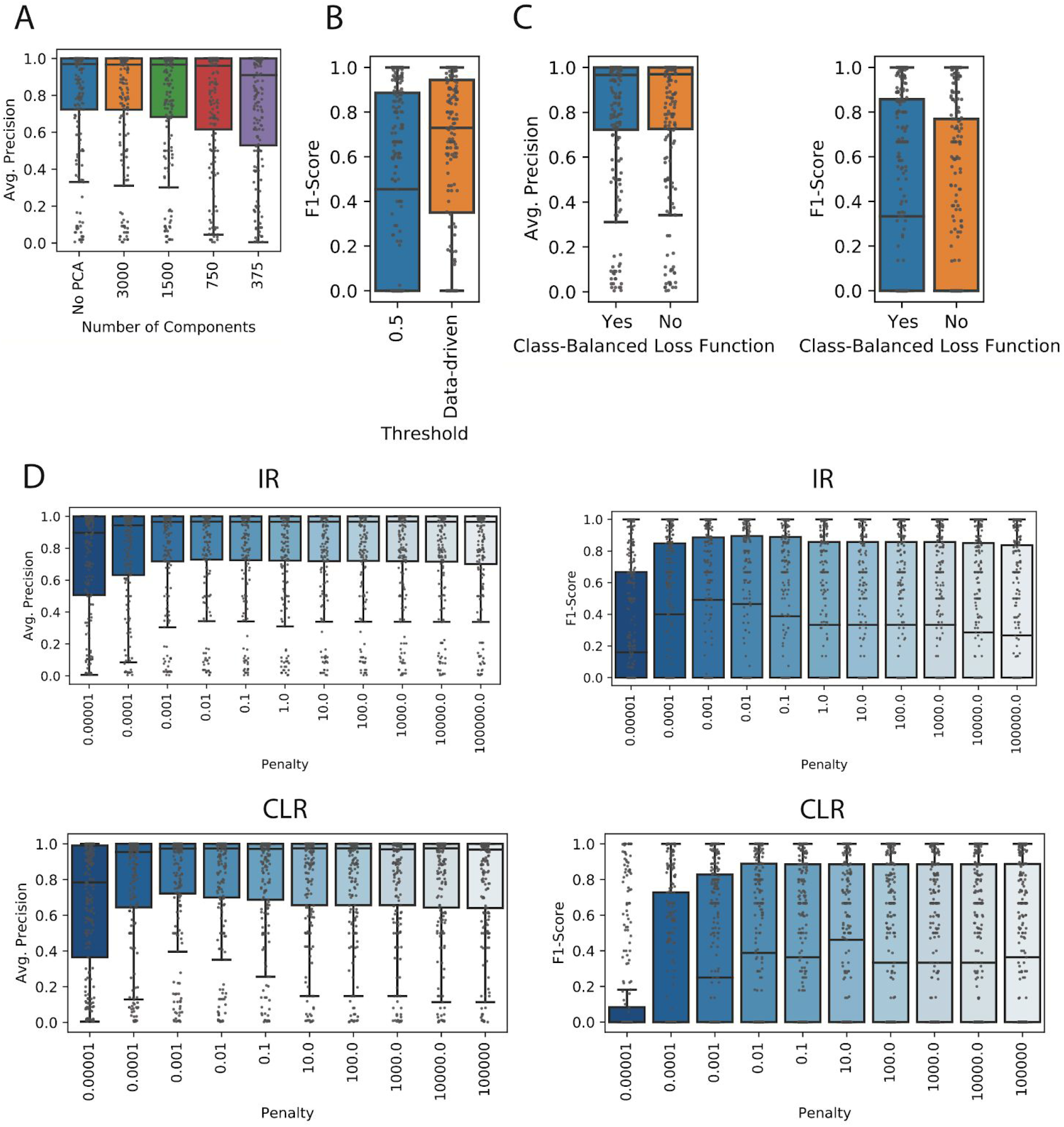
Parameter tuning. (**A**) Average precision scores across cell types in the bulk RNA-seq validation set produced by the independent one-vs.-rest classifiers on data preprocessed with various numbers of principal components. (**B**) F1-scores produced by the independent classifiers when using a threshold of 0.5 for making a binary yes-no decision for all cell types versus a custom threshold for each cell type as empirically determined via a leave-study-out cross-validation experiment on the pre-training set of bulk RNA-seq samples. (**C**) Average precision (left) and F1-scores (right) across cell types in the bulk RNA-seq validation set produced by the independent one-vs.-rest classifiers trained either with or without the class-balanced loss-function. (**D**) Average precision scores across cell types in the bulk RNA-seq validation set produced by the one-vs.-rest and CLR conditional classifiers when trained with various regularization strengths (i.e. penalties). These penalty values correspond to the inverse of the regularization strength and thus, smaller numbers indicate stronger regularization.

**Supplementa1 Figure 6.**
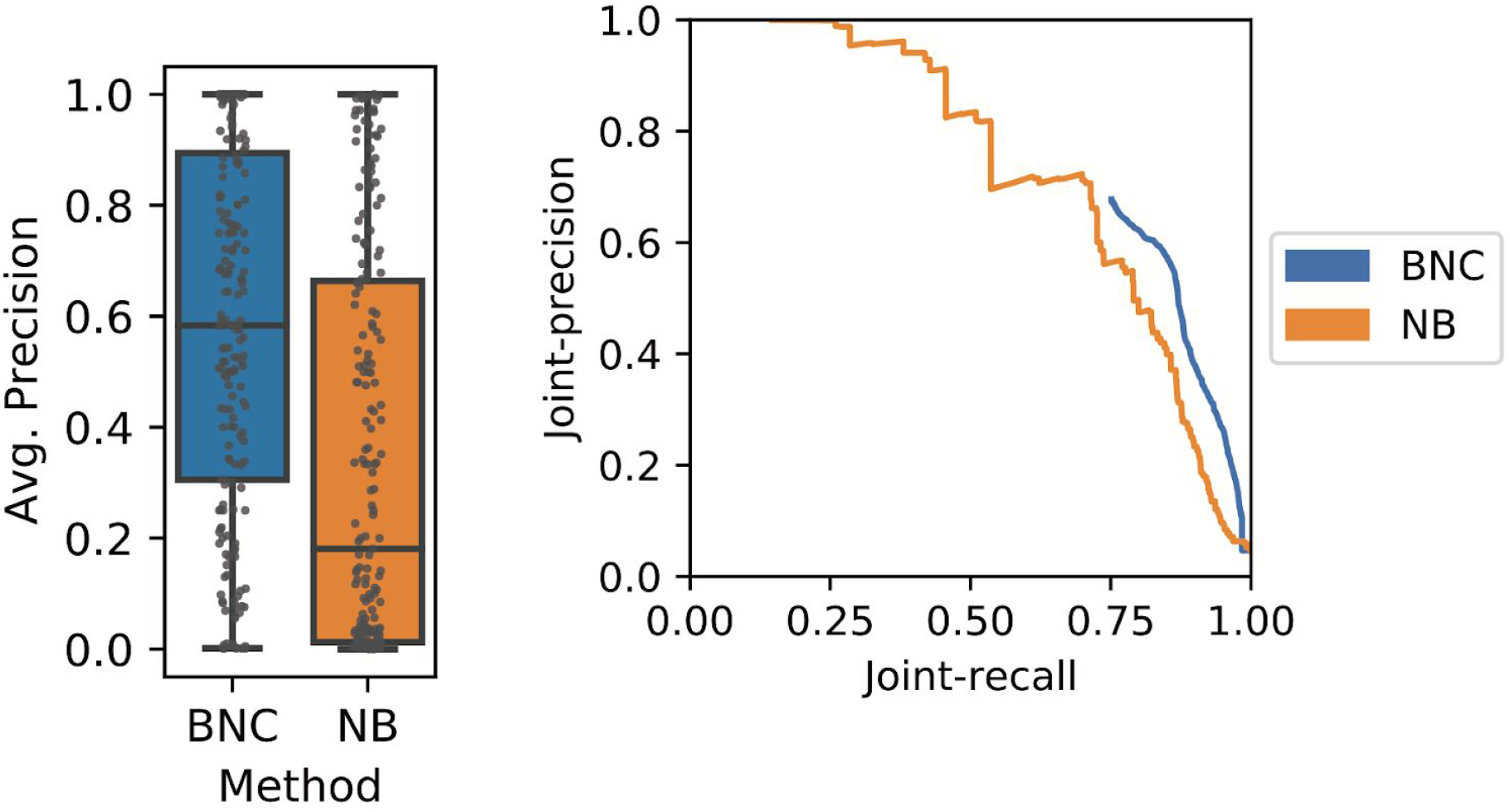
Comparing Bayesian network correction with naive Bayes. A comparison between BNC and a naive Bayes variant of BNC that considers each cell type independent of the graph-dependencies between cell types. We compared these approaches in terms of both the distribution of average precisions across cell types (left) and joint precision-recall curves constructed by considering each cell-cell type pair as an independent prediction.

**Supplementa1 Figure 7.**
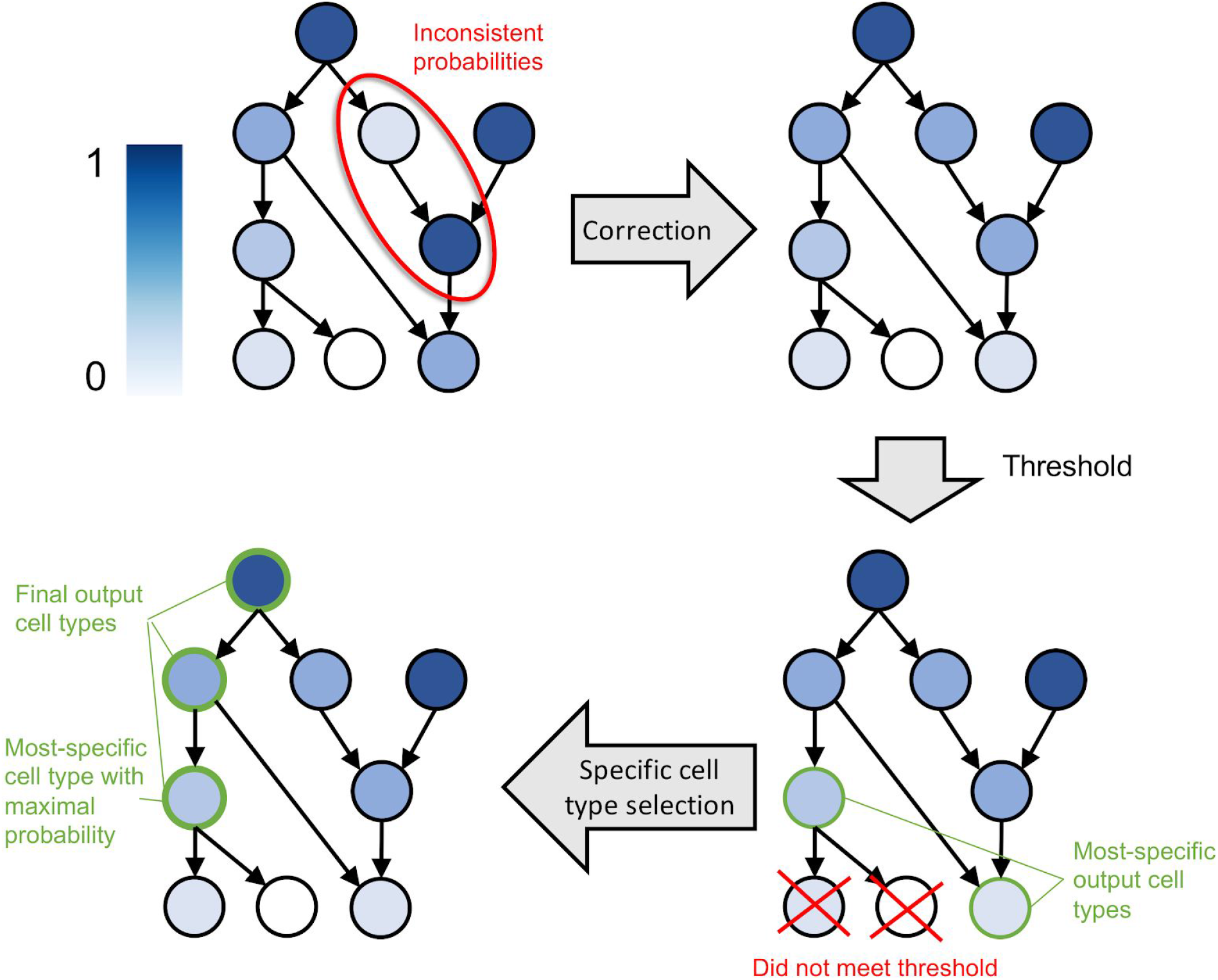
Binarizing CellO’s classifier probabilities. A schematic illustration of how CellO outputs binary yes-no cell type classifications for a given sample. First, the raw probabilities are corrected with the cell ontology using IR (if CLR is used, this step is not necessary). Cell types whose raw probabilities meet their respective decision-threshold are selected. Among these, the most-specific cell types (i.e. lowest in the ontology) are examined and the cell type with the highest output probability is selected. CellO outputs this final selected cell type along with all ancestor terms.

**Supplementa1 Figure 8.**
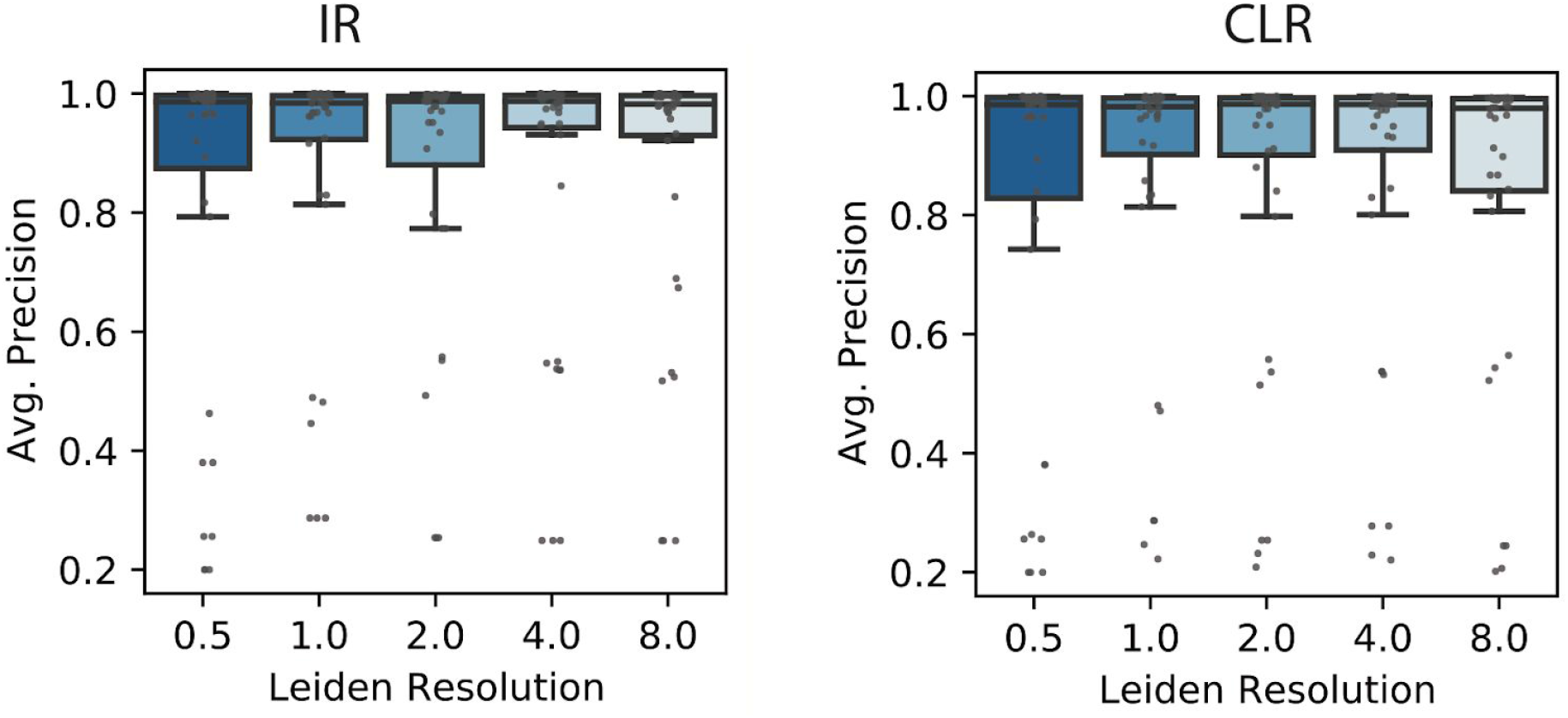
Effect of clustering parameter on performance. We clustered the Zheng et al. (2017) PBMC dataset using various resolution parameters for the Leiden community-detection algorithm and then computed the average-precision across all cell types produced by IR (left) and CLR (right) on the mean expression profiles for the clusters generated under each parameter.

**Supplementa1 Figure 9.**
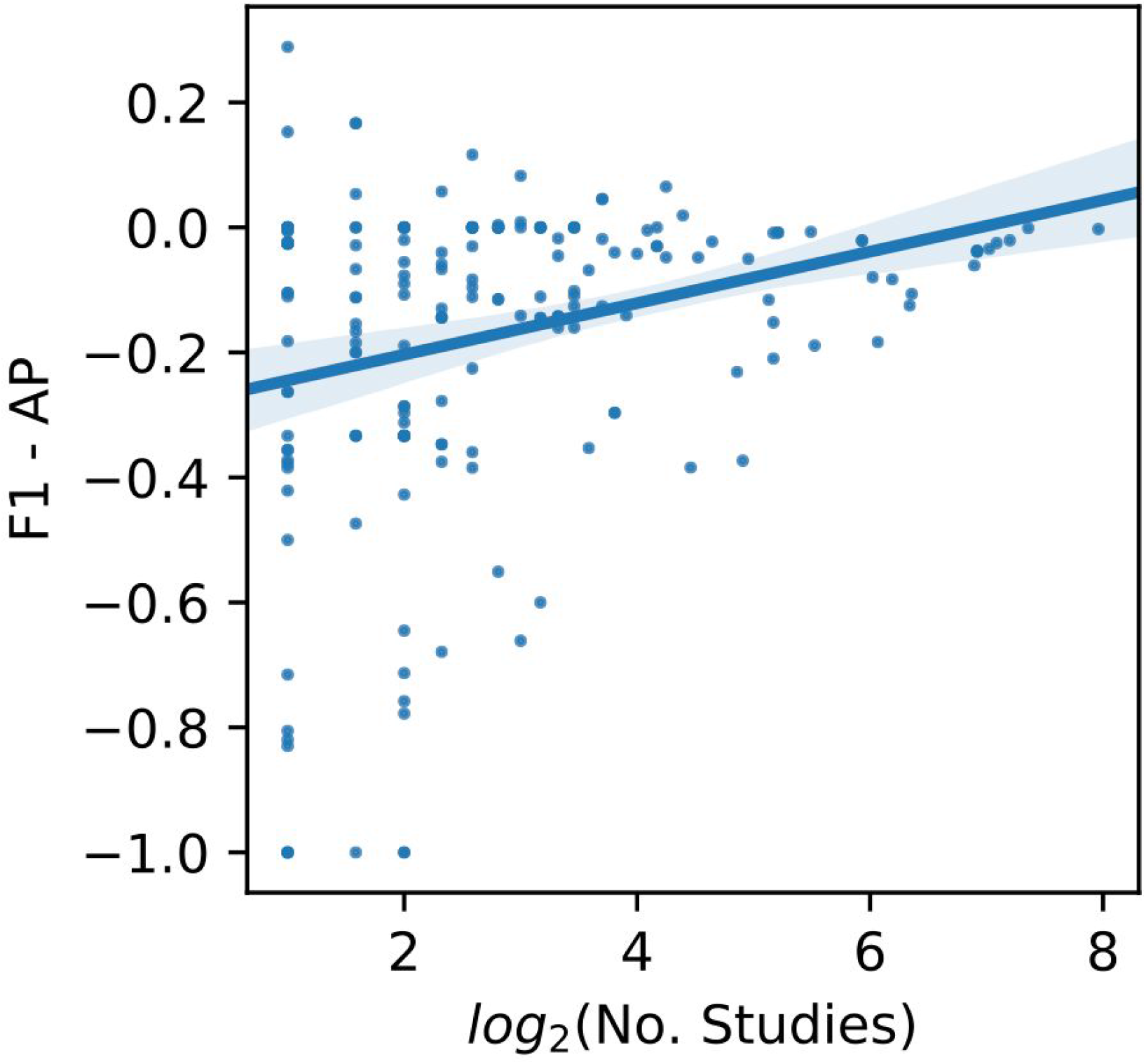
Effect of number of training studies on calibration quality. We evaluated IR on the bulk RNA-seq validation set and for each cell type, we computed the F1-score (F1) using a decision-threshold of 0.5 and average precision (AP). We then plot the difference between F1 and AP against the logarithm of the number of studies in the bulk RNA-seq pre-training set that sequenced each cell type. A larger gap between F1 and AP (i.e. negative numbers with large magnitude) indicates poorer calibration. Error bands around the ordinary least-squares regression line indicate 0.95 confidence intervals estimated via bootstrapping.

